# Tissue surface mechanics constrains proliferation-driven forces to guide mammalian body axis elongation

**DOI:** 10.1101/2025.10.27.684710

**Authors:** Marc Trani-Bustos, Ryan G Savill, Arthur Boutillon, Petr Pospíšil, Deniz Conkar, Claudia Froeb, Johannes R Soltwedel, Heidi L van de Wouw, Ellen M Sletten, Jesse V Veenvliet, Otger Campàs

## Abstract

Mammalian embryos undergo complex morphogenetic changes after implantation in the uterus. The elongation of the body along a head-to-tail axis is a pivotal event, as it lays the foundation of the body plan. While genetic and biochemical aspects of mammalian body elongation have been uncovered, the physical mechanism of axial morphogenesis remains unknown, largely due to the inaccessibility of the implanted embryo to physical measurements and manipulations *in utero*. Gastruloids, a stem-cell-based embryo model of mammalian axial morphogenesis, lift such limitations. Combining live imaging, direct mechanical measurements, and chemical and mechanical perturbations, here we show that axis elongation in mouse and human gastruloids is guided by a posterior ‘actin cap’ at the tissue surface that constrains the expansive forces of cell proliferation. Measurements of mechanical stresses using oil microdroplets, as well as inhibition of cell proliferation and myosin activity, show that the forces driving elongation arise from cell proliferation, and not from convergent extension movements. We find that isotropic tissue expansion is redirected into posterior elongation by the formation of a supracellular actin cap at the posterior tissue surface that restricts lateral tissue expansion. Finally, we show that posterior elongation in mouse embryos displays the key features of the physical elongation mechanism reported for mouse and human gastruloids. These findings reveal that mammalian body axis elongation, including human, occurs via a different physical mechanism from other vertebrate species.

---

Early embryogenesis involves dynamic shape changes that result in the establishment of the body axes — the coordinate system for future organ formation. One of the most universal characteristics of embryogenesis is directional elongation of the early embryo along the head-to-tail, or anteroposterior (AP), body axis^1,2^ . Recent studies have revealed how mechanics shapes the elongating body axis in chick, amphibians and fish, and point toward distinct physical mechanisms of elongation in different species^3,4^. While in some cases, such as amphibians, convergent extension is essential for elongation^5,6^, in chick or fish the existence of material (fluid/solid) transitions along the AP axis plays a key role in elongation^7,8^. It is unclear whether mammalian species, including humans, physically elongate their AP axis using mechanisms similar to other vertebrate species, or instead evolved a different physical mechanism of axis formation.

Direct mechanical measurements of mammalian embryos *in vivo* are very challenging, since the embryo develops within the uterine wall^9,10^. Stem-cell-based embryo models, in which pluripotent stem cells self-organize into embryo-like structures, provide an opportunity to study morphogenesis in mammals^10–13^ . To elucidate the physical mechanism of mammalian body axis elongation, we took advantage of gastruloids, a stem-cell-based embryo model that recapitulates key features of the natural embryo^10,12,14,15^ . Upon treatment of a spherical stem cell aggregate with a WNT agonist, gastruloids break symmetry and elongate along the AP axis between 96h and 120h after aggregation, expressing — like in the embryo — the transcription factor Brachyury (Bra) at the posterior end^10,12,14,16–19^ (Fig. 1a,b). To reveal the physical mechanism of elongation in mammals, we combined mechanical measurements and perturbations in both mouse and human gastruloids, and verified in mouse embryos that the key features of the physical mechanism discovered in gastruloids are also observed *in vivo*.

**Figure 1.**
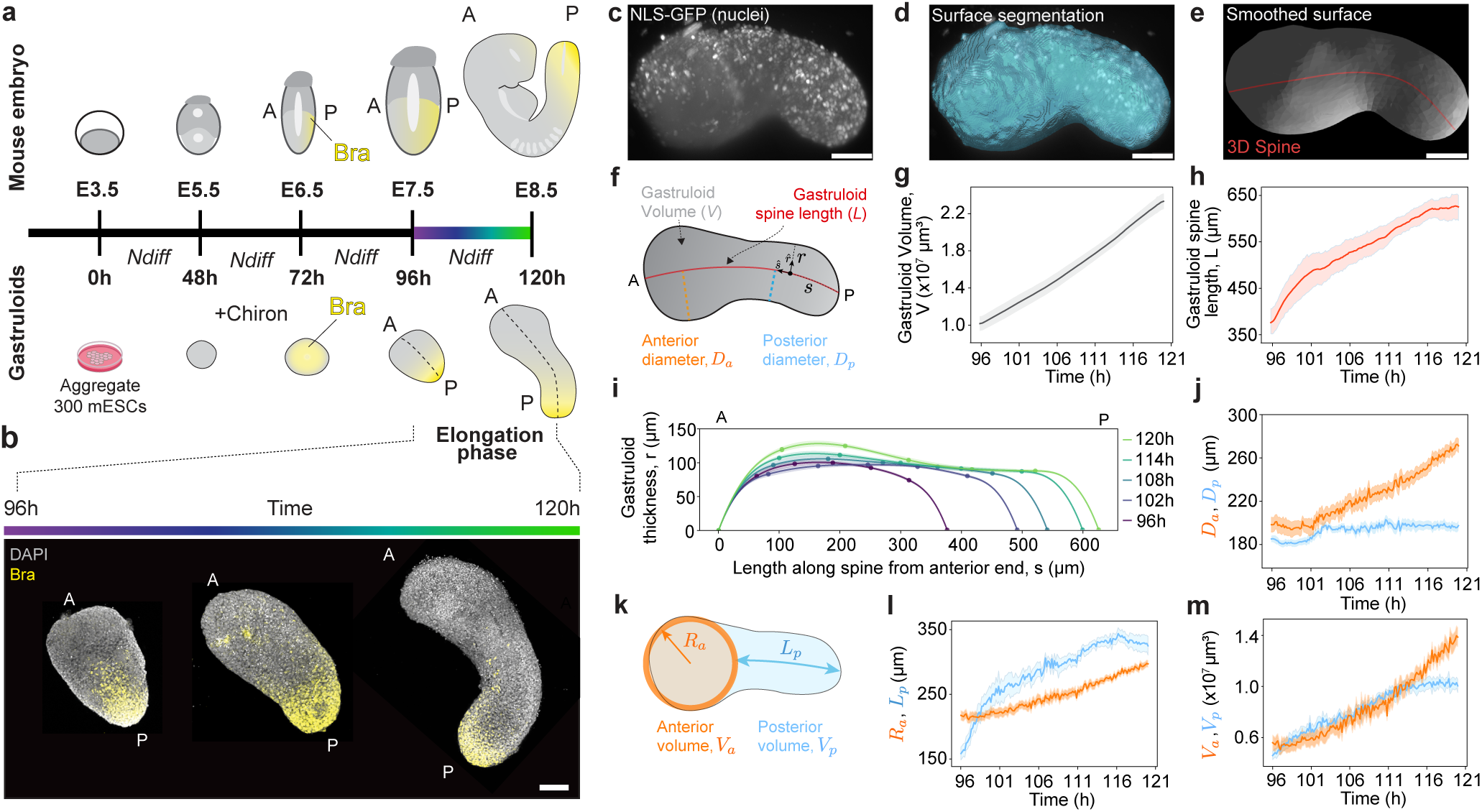
Morphodynamics of mouse gastruloid elongation. **a,** Schematic representation of a developing mouse embryo (top) and a mouse gastruloid at the equivalent stages (bottom). Yellow, Brachyury (T/Bra, posterior marker); E, embryonic day; A, anterior; P, posterior; NDiff, gastruloid differentiation medium. Purple-to-green color gradient indicates time during the gastruloid elongation phase (96h – 120h). **b,** Representative Maximum Intensity Projections (MIPs) of nuclei (DAPI; gray) and Bra (yellow) in fixed mouse gastruloids at different stages of axial elongation. **c,** Representative maximum intensity projection (MIP) of a nuclear-labeled (NLS-GFP) gastruloid at 120h. **d-e,** 3D surface segmentation (**d**), surface mesh smoothing and 3D spine extraction (**e**) of the gastruloid shown in **c**. **f,** Sketch of a gastruloid indicating the local coordinate system along its spine (*s* and *r* being the spine contour length from posterior end and the local gastruloid thickness; *s*^, *r*^ being the associated local tangential and radial unit vectors), its total volume V (gray), spine length L (red), as well as anterior (orange) and posterior (blue) diameters, *D_a_* and *D_p_* respectively. **g-h,** Gastruloid volume (**g**) and spine length (**h**) during elongation. Mean (line) and s.e.m (band) are shown. **i,** Time evolution of the axisymmetric gastruloid shape (ensemble average); measured average thickness (dots) and cubic spline (lines). Color code as in **a**. **j**, Measured anterior and posterior diameters, *D_a_* and *D_p_*, during elongation. **k,** Scheme of the spherical anterior domain (orange) and posterior domain (blue), characterized by their anterior radius *R_a_* and posterior length *L_p_*, as well as their volumes *V_a_* and *V_p_* , respectively. **l-m,** Time evolution of geometrical characteristics (**l**) and volume (**m**) of anterior and posterior domains. Color code as in **k**. N=12 gastruloids in **g**-**h**, **i**, **l**-**m**. Scale bars, 100 μm.

## Mouse gastruloids elongate posteriorly as cylinders of constant diameter

To characterize the morphogenetic changes during gastruloid elongation, we performed light-sheet 3D live imaging of gastruloids generated from mouse embryonic stem cells (mESCs) with fluorescently labelled nuclei (NLS-GFP^20^; Fig. 1c; Extended Fig. 1a-d; Methods). Using a computational framework that we recently developed (SpinePy^20^ ; Methods), we monitored the 3D gastruloid geometry over time by segmenting its surface and calculating its spine (Fig. 1d-f; Extended Fig. 1e; Supplementary Video 1), which defines the intrinsic coordinate system for each gastruloid and enables ensemble statistics across gastruloids. Both the gastruloid volume V and AP length L (spine length) increased steadily between 96-120h, only tapering off after 116h (Fig. 1g,h). Analysis of the axisymmetric gastruloid shape (Fig. 1i; Extended Fig. 1f-h; Methods) showed that its posterior half maintains a constant diameter during elongation (Fig. 1j), resembling a cylinder, while the anterior end expands isotropically, resembling a sphere. Both the radius of the anterior spherical domain (Fig. 1k) and the length of the posterior domain increased during gastruloid growth (Fig. 1l). The observed volumetric growth of the gastruloid contributes both to elongate the posterior cylindrical domain and expand the anterior spherical domain (Fig. 1m), up to about 116h, when posterior elongation stops and the anterior domain keeps expanding. These data show that gastruloids grow by both the isotropic expansion of an anterior spherical domain and concomitant elongation of a cylindrical posterior domain.

## Cell proliferation causes the volumetric growth necessary for elongation

Since gastruloid elongation involves volumetric growth, we investigated whether cell proliferation was necessary for elongation. Cell division events were uniformly distributed throughout the gastruloid at all timepoints (Fig. 2a), as indicated by their constant 3D density along the AP axis (Fig. 2b-c; Methods). While the total number of cell division events increased over time from 96-110h and stagnated after that, the density of cell division events was maintained constant due to the proportional increase in gastruloid volume (Fig. 2d-e), revealing a constant cell division density in both space and time during gastruloid growth. Inhibition of cell proliferation from the onset of the elongation phase using Genistein, which arrests the cell cycle at G2-M^21^, led to a dose-dependent decrease of cell division events (Fig. 2f,g) without causing substantial cell death or affecting Bra expression (Extended Fig. 2). Similar effects were observed upon treatment with other cell cycle inhibitors (Extended Fig. 3). Neither volume growth (Fig. 2h) nor elongation (aspect ratio; Fig. 2i) were observed above 10 µM Genistein concentration, coinciding with a substantial suppression of cell division events (Fig. 2g). Removal of Genistein after 6h of treatment (wash-out) showed that its effect is reversible, with the gastruloid resuming elongation and recovering in volume and aspect ratio (Fig. 2h-i). These data show that cell proliferation is essential for gastruloid growth and its posterior elongation.

**Figure 2.**
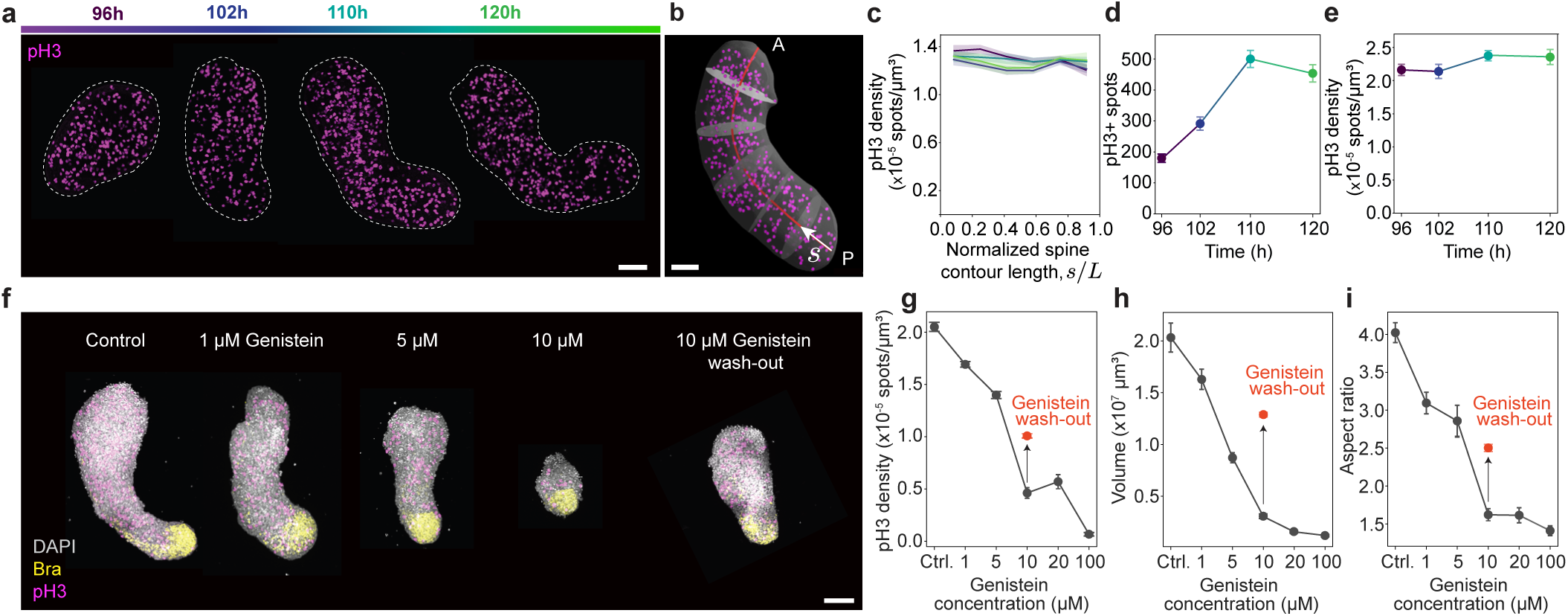
Cell proliferation is essential for both growth and elongation of mouse gastruloids. **a,** Mouse gastruloids (MIPs) during elongation (96h, 102h, 110h, 120h) stained for mitotic marker phospho-histone3 (pH3, magenta). Time color code as in Fig. 1a. **b,** Representative 3D view of pH3 in a gastruloid (110h; transparent surface segmentation shown), with its spine (red) and disks (gray) defining different regions along the AP contour. **c,** Measured pH3+ nuclei (cell division events) density along the AP axis. Different lines correspond to different timepoints. **d-e,** Temporal evolution of the absolute number of pH3+ nuclei and pH3 density (absolute number/total volume). Color code as in **a**. N= 76, 60, 46, 23 gastruloids for 96h, 102h, 110h and 120h samples, respectively, for **c**-**e**. **f,** Confocal images (MIP) of fixed gastruloids stained for nuclei (DAPI, gray), T/Bra (yellow) and pH3 (magenta) in control conditions and in samples treated with different Genistein concentrations to inhibit cell divisions. A wash-out condition in which Genistein (10 μM) was removed and gastruloids allowed to recover until 120h is also shown. **g-i**, Measured pH3 density (**g**), as well as gastruloid volume (**h**) and aspect ratio (**i**) for increasing Genistein concentration (dark gray). Values for the Genistein wash-out condition are also shown (red; arrow indicates recovery). N = 15, 25, 20, 13, 11, 12 gastruloids for control, 1, 5, 10, 20, 100 μM Genistein conditions, respectively. N= 23 gastruloids for Genistein wash-out. Mean (dots) and s.e.m. (error bars) are shown. Scale bars, 100 μm.

## Proliferation-driven mechanical stresses, but not convergent extension, drive gastruloid elongation

Despite the relevance of cell proliferation for volumetric growth, other processes, such as convergent extension (CE), are known to be important for axis elongation in several species^3,6,22^. To understand the origin of the forces driving elongation, we used oil microdroplets^7,23^ to directly quantify the orientation and magnitude of the mechanical stresses in elongating gastruloids (Methods). We first adapted the microdroplet technique for gastruloids by introducing custom fluorosurfactants^24^ that substantially lowered the droplet interfacial tension, enabling reliable measurements of the small stresses in gastruloids (Methods). After injecting a single, calibrated droplet in the gastruloid, we imaged and reconstructed the droplet in 3D during gastruloid elongation (Fig. 3a-b; Methods). Using the STRESS software^25^, we obtained the magnitude and principal axes of tissue-scale anisotropic stresses from droplet deformations (ellipsoidal deformation mode; Fig. 3c; Methods). The angle between the direction of droplet elongation (main principal axis) and the AP axis provides information about the origin of the stresses^7,26^. If actomyosin-generated active stress anisotropy drives elongation (dominant stress along the mediolateral direction), as in CE, droplets align along the AP axis (Fig. 3d). Instead, droplets align in the perpendicular direction if they resist some other internal stresses, such as those generated by proliferation^26^ (Fig. 3d). Analysis of droplet orientation revealed that droplets elongate perpendicular (approx. 90 degrees) to the local direction of the AP axis at all timepoints (Fig. 3e; Methods). Deviations were only observed at 120h, for which droplet orientations preferentially displayed angles between 60-70 degrees (Fig. 3e). These results indicate that the mechanical stresses driving elongation do not originate from a process of CE, in agreement with the observation that the gastruloid does not become thinner during elongation (Fig. 1i-j). Indeed, pharmacological inhibition of non-muscle myosin II using blebbistatin led to increased elongation, as indicated by the larger aspect ratio of the resulting gastruloids (Fig. 3f-g), in direct contrast with what is expected in CE.

**Figure 3.**
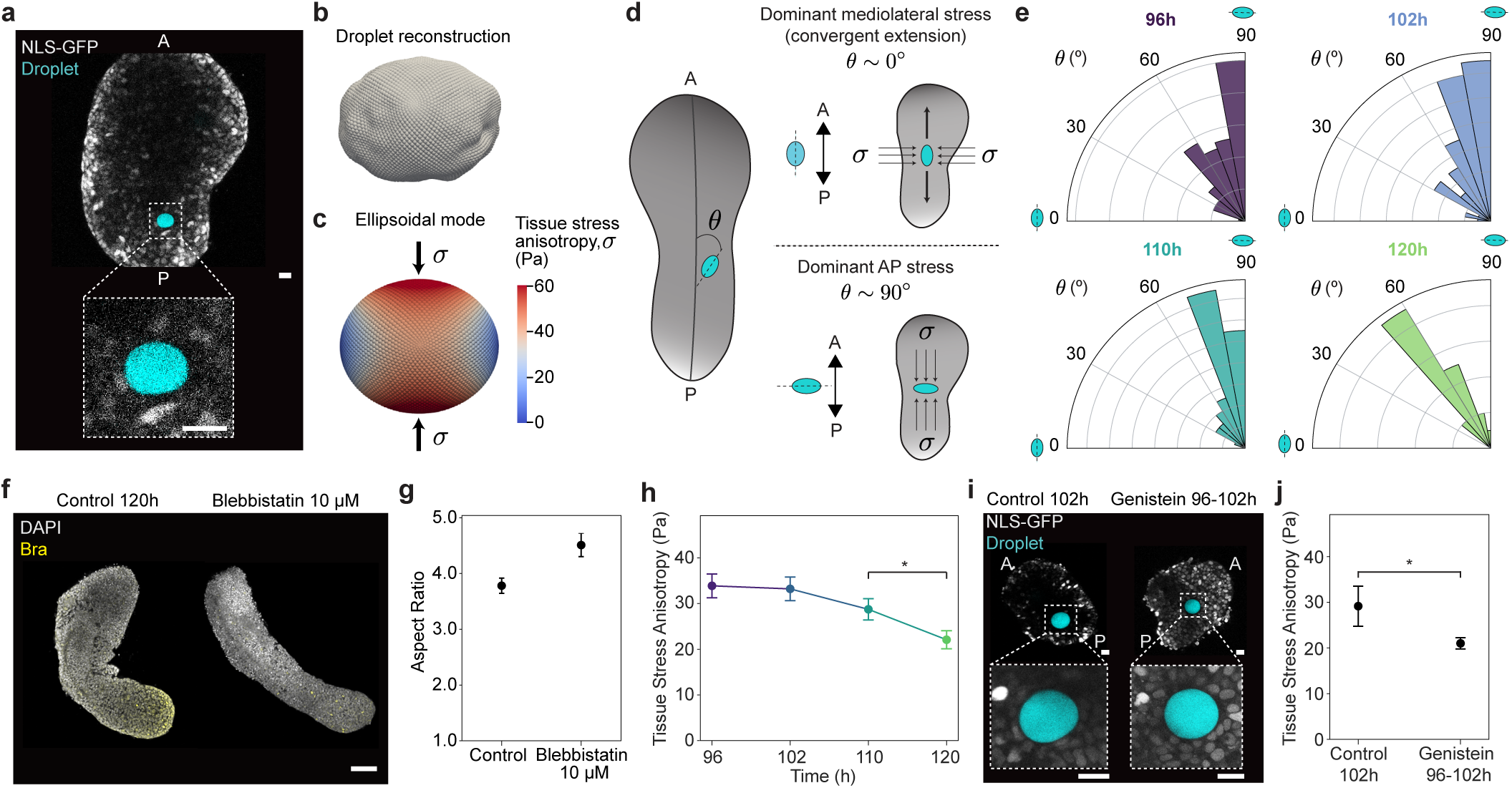
Mechanical stresses driving gastruloid elongation. **a**, Live imaging (confocal section) of a nuclear labeled (NLS-GFP, gray) gastruloid with a previously inserted oil droplet (cyan). The dashed square shows a magnified region with the oil droplet. Scale bars, 20 μm. **b-c**, 3D reconstruction of the droplet surface (**b**; gray) and associated ellipsoidal deformation mode with the surface mean curvature values mapped onto the surface (**c**). Arrows in **c** indicate the stresses *σ* applied on the droplet. **d**, Sketch defining the angle *8* between the AP axis and the long axis of the droplet’s ellipsoidal mode, as well as the expected angles for specific limiting scenarios: dominant mediolateral stresses (e.g., convergent extension) or dominant AP oriented stresses. **e**, Measured values of the angle *8* in gastruloids at 96h, 102h, 110h and 120h (time color code as in Fig. 1a). n (N) = 82 (31), 48 (16), 42 (16), 19 (5) droplet measurements (gastruloids) for 96h, 102h, 110h and 120h, respectively. **f**, Confocal imaging (MIPs) of fixed gastruloids (120h) in control conditions and in Blebbistatin 10μM to inhibit non-muscle myosin II. Scale bar, 100 μm. **g**, Measured gastruloid aspect ratio in control and in Blebbistatin-treated gastruloids. N= 25, 22, respectively. **h**, Time evolution of tissue-scale stress anisotropy in elongating gastruloids. p = 0.033 (*; Welch’s test). n (N) = 82 (31), 54 (17), 47 (18) and 27 (9) droplet measurements (gastruloids) per time point, respectively. **i**-**j**, Droplet stress measurements in gastruloids with inhibited cell proliferation. Confocal sections of control (left) and Genistein (10μM) for 6h (right) gastruloids (**i**), and associated measurements of tissue-scale stresses (**j**). p = 0.023 (*; Mann-Whitney U test). Scale bars, 20 μm. N= 28, 38, respectively. Mean (dots) and s.e.m. (error bars) shown.

To test whether cell proliferation was causing the mechanical stresses necessary for elongation, we measured the magnitude of tissue stress anisotropy both in the presence and absence of cell proliferation. Tissue stresses were uniform along the AP axis at all timepoints during normal elongation (Extended Fig. 4a), mirroring the uniform and constant proliferation density (Fig. 2a-e). Their magnitude was largely constant over time during elongation, with only a slight but significant decrease at 120h (Fig. 3h), coinciding with the observed slowdown in gastruloid elongation (Fig. 1h). Finally, inhibiting cell proliferation led to a significant decrease in mechanical stress (Fig. 3i-j). Together, these data show that CE does not elongate mouse gastruloids, and instead cell proliferation is the driving force behind elongation.

## A posterior actin cap constrains lateral tissue expansion, guiding posterior elongation

While proliferation-driven mechanical stresses fuel gastruloid elongation, how these forces translate into unidirectional elongation remains unclear. Given the role of cell proliferation, oriented cell division events could be at the origin of anisotropic tissue growth^3,27^. To examine a potential role of oriented cell division, we measured the direction of cell division with respect to the local direction of the AP axis (Fig. 4a). Cell divisions appeared randomly oriented during elongation, as revealed by the angular distribution of the mitotic planes of pH3 positive cells between 96-120h (Fig. 4a-b). These results show that gastruloid elongation is not caused by oriented cell divisions.

**Figure 4.**
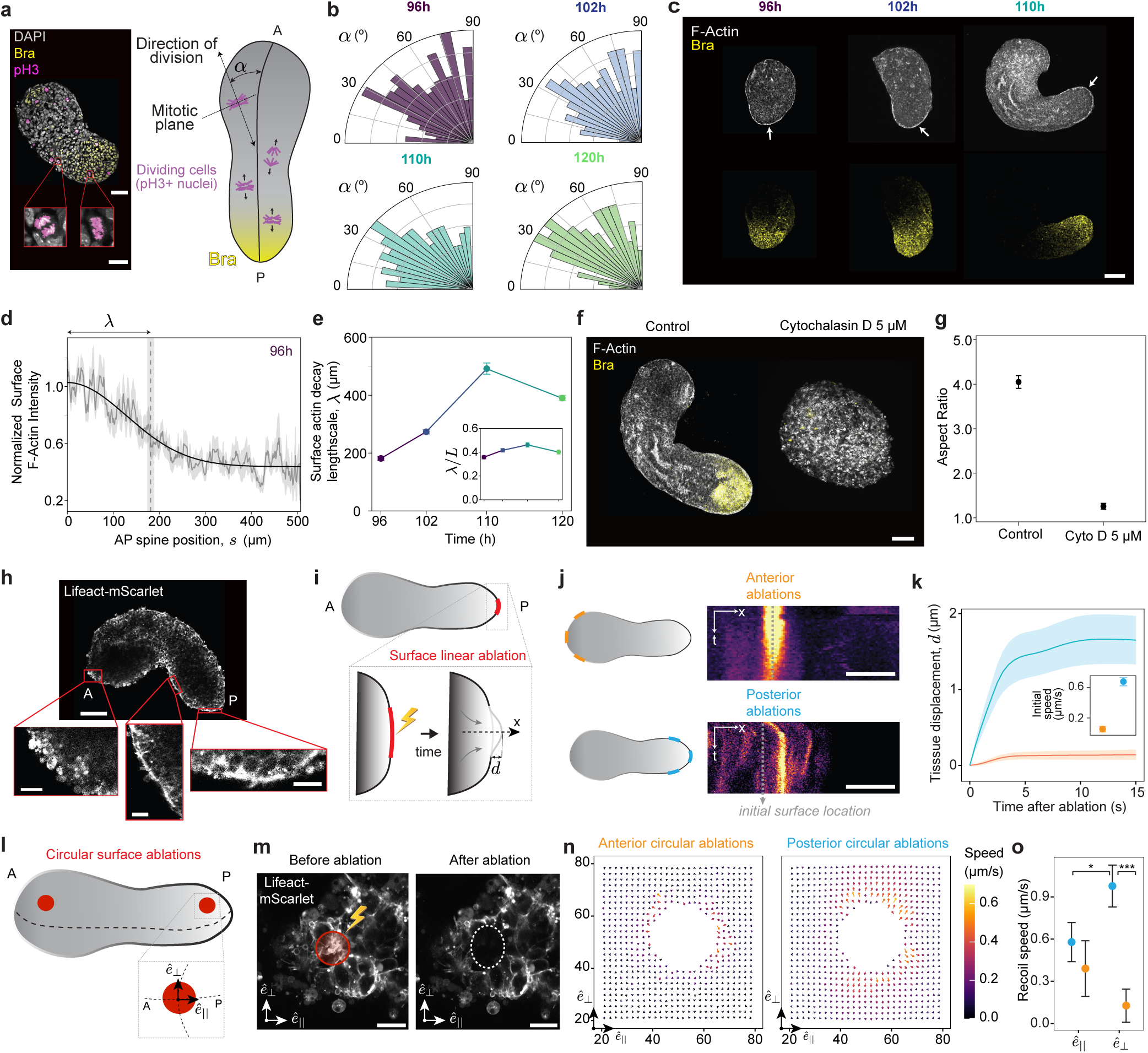
An actin cap restricts lateral gastruloid expansion and guides elongation. **a**, Confocal section of a fixed gastruloid (left) stained for nuclei (DAPI, gray), dividing cells (pH3, magenta), and Bra (yellow). Examples of dividing cells (red squares; anaphase, left; metaphase, right). Sketch defining the angle *α* between the direction of cell division and the local direction of the AP axis. **b**, Measured orientation of cell division during elongation (96h, 102h, 110h, 120h). n (N) = 401 (10), 390 (10), 226 (5), 131 (3) cells (gastruloids) for each time point, respectively. Time color code as in Fig. 1a. **c**, Confocal imaging (MIP of 20-30 equatorial planes) of fixed mouse gastruloids stained for F-Actin (Phalloidin) and Bra (96h, 102h, 110h). Arrows indicated the posterior high intensity of surface actin. Scale bar, 100 μm. **d**, Surface actin intensity along the AP axis at 96h (mean, gray line; sem, gray band; fit, black line). The decay length scale *λ* is shown (vertical dashed line). **e**, Measured *λ* during elongation (96-120h; mean ± sem), and value normalized to the gastruloid length *L* (inset). N= 11 (96h), 11 (102h), 12 (110h), 12 (120h) measured gastruloids. **f-g**, Confocal images (MIP) of fixed gastruloids (120h) stained for F-Actin (Phalloidin) and Bra in control conditions and in 5 μM Cytochalasin D (**f**; scale bar, 100 μm), and measured gastruloid aspect ratio in each condition (**g**; N = 13, 12, respectively). **h**, Confocal live imaging of an actin-labelled gastruloid (Lifeact-mScarlet, gray; 115h), and closeups of surface actin organization at anterior (left), middle (center), and posterior (right) regions. Scale bar, 100 μm for overview, and 20 μm for closeups. **i**, Sketch of laser ablation (red patch) of actin at the gastruloid surface and the resulting tissue expansion, *d*, in the normal direction, *x*. **j-k**, Kymographs of the time evolution of actin intensity after ablation along the normal direction at anterior (orange) and posterior (blue) regions (**j**; sketches in left show regions of ablation; scale bars, 10 μm), along with quantification of tissue expansion in anterior and posterior regions (**k**; initial expansion speed, inset). N = 40 rectangular ablations; 10 gastruloids. **l-o,** Sketch showing circular surface ablations in posterior and anterior patches (**l**; red; local coordinate system defined in magnified region). Confocal images of circular surface actin (Lifeact-mScarlet) ablation before and after ablation (**m;** scale bars, 10 μm) along with associated PIV quantifications of surface tissue movements (**n**), and measured recoil speed of tissues along the AP axis and circumferential direction (**o**). Mann-Whitney U test with p = 0.022 (*) and p = 0.0001 (***) indicated. N = 30 circular ablations; 10 gastruloids.

Beyond anisotropy in bulk tissue growth, unidirectional elongation could be guided by mechanical constraints at the surface of the gastruloid. Inspection of the actin distribution at the gastruloid surface showed an enrichment of actin at its posterior domain from 96-120h (Fig. 4c). Quantification of the surface actin signal along the AP axis showed a higher signal at the posterior end that decayed toward the anterior end over a length scale *λ* of approximately half the gastruloid length (Fig. 4d; Methods), creating a posterior actin cap that persists for all timepoints during extension (96-120h; Extended Fig. 5). The length *λ* of the actin cap increased from 96-120h following elongation (Fig. 4e), always covering the posterior domain that spans to about half the gastruloid length (Fig. 4e, inset). We hypothesized that this posterior actin cap could laterally constrain the proliferation-driven stresses to guide elongation. Disruption of the actin cap with Cytochalasin D completely suppressed gastruloid elongation, as indicated by a substantial decrease of their aspect ratio (Fig. 4f-g), but not their growth, leading to aggregates that grew isotropically (Extended Fig. 6). These results suggest that the posterior actin cap acts as a surface mechanical constraint that restricts lateral expansion and guides gastruloid elongation.

To enable real-time visualization and manipulation of the actin cap, we engineered mESCs with LifeAct-mScarlet and generated gastruloids (Methods). The surface LifeAct signal revealed a smooth, flat supracellular actin structure at the posterior domain (Fig. 4h), consistent with a mechanical constraining role. In contrast, the anterior domain lacked such organization, showing disorganized actin and rounded cells at the surface (Fig. 4h). To test whether the posterior actin cap could mechanically constrain the expanding tissue in its interior, we performed laser ablation of the surface actin and monitored the resulting movement of the underlying bulk tissue along the normal direction to the surface (Fig. 4i-k; Supplementary Videos 2-3). In the gastruloid posterior domain, the tissue expands outwards following local ablation of the actin cap (Fig. 4j-k). In contrast, ablation of surface actin in the anterior domain leads to no outward expansion of the tissue underneath (Fig. 4j-k). These data indicate that the posterior actin cap is resisting the expansion of the tissue.

To further test the response of surface actin to mechanical perturbations, we performed surface circular ablations in both anterior and posterior gastruloid domains (Fig. 4l-m; Supplementary Videos 4-5), and monitored the surface displacement field. While barely any surface movement was observed in response to surface ablation at the anterior domain (Fig. 4n-o), circular ablation of the actin cap in the posterior domain caused an anisotropic recoil of the surface. Maximal recoil at the posterior domain occurred along the circumferential direction (perpendicular to the AP axis; Fig. 4l,o), as expected for the tubular geometry of the actin cap. Altogether, these results show that a posterior actin cap, which covers the posterior domain as it elongates, constrains the internal expansive forces caused by cell proliferation, thereby guiding AP elongation.

## Posterior elongation in mouse embryos displays the same key features as mouse gastruloids

To understand whether the same physical mechanism of elongation is at play in mouse embryos, we tested whether the two key features of this mechanism, namely homogeneous cell divisions in the elongating tissues and an actin cap at their surface, are present during posterior elongation *in vivo* (Fig. 5a). Since mouse embryos are considerably more complex than gastruloids due to the presence of multiple tissues interacting with each other, we focused on the presomitic mesoderm (PSM) as it is the tissue that elongates the body axis in many species^2,3^ and is mirrored by gastruloid elongation^10^ (Fig. 5b). As for mouse gastruloids, cell divisions were present and uniform from the tailbud to the PSM of E8.5 mouse embryos (Fig. 5b-c). Inhibiting cell proliferation in tail explants strongly suppressed cell divisions (Fig. 5d-f) and arrested posterior elongation (Fig. 5g; Supplementary Videos 6-7), in agreement with our results in gastruloids. Quantification of actin intensity at the surface of the tissue along the the AP axis (tailbud-PSM-somites) showed a posterior-to-anterior gradient (Fig. 5h-i) similar to that observed for gastruloids (Fig. 4c,d). These results show that the key features of the physical elongation mechanism reported above for mouse gastruloids are present *in vivo*, suggesting that posterior body elongation in mouse embryos shares the same physical mechanism of axis elongation as mouse gastruloids.

**Figure 5.**
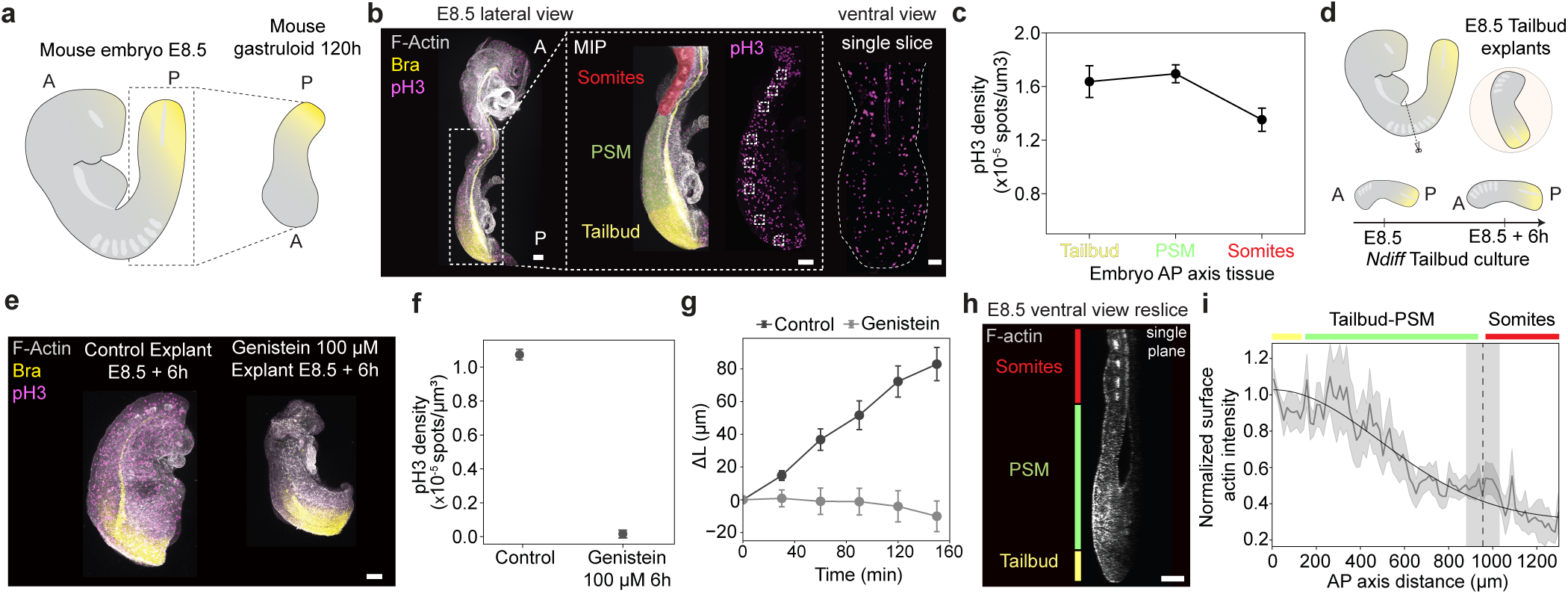
Homogeneous cell divisions and a posterior actin cap are present in elongating mouse embryos. **a**, Sketch comparing elongating mouse embryos at E8.5 (left) and mouse gastruloids at 120h, with Bra expression as posterior marker (yellow). **b**, Lateral confocal image (MIP) of a fixed E8.5 mouse embryo stained for F-Actin, Bra and pH3 (left). Magnified posterior trunk region (dashed square) colored by tissue region (tailbud, yellow; PSM, green; somites, red). pH3 signal is shown separately in lateral view (MIP) and ventral view. Small dashed squares in lateral view indicate regions of pH3 density measurements along the AP axis. **c**, Measured pH3 density along the A-P axis in the tailbud, PSM and somite regions of an E8.5 mouse embryo. N= 22, 25, 30 per region, respectively; n=11 embryos. **d**, Sketch showing a mouse embryo E8.5 tailbud explant experiment, where mouse embryo tails are manually dissected from the 4th last somite and cultured with Ndiff medium for 6h in control or in 100 μM Genistein. **e**, Confocal images (MIPs) of fixed mouse tail explants stained for F-Actin, Bra and pH3 after 6h culture in control and in 100 μM Genistein conditions. Scale bar, 100 μm. **f-g,** Measured pH3 density (**f**) and relative tail length change Δ*L* (**g**) in control and in Genistein-treated mouse embryo tailbud explants. N= 11, 14, respectively. **h**, Ventral view reslice along the tailbud-somites regions of a confocal 3D image of a fixed E8.5 mouse embryo stained for F-Actin (gray), showing a decrease of surface actin intensity from the tailbud (yellow) and PSM (green) to the somites (red). **i**, Measured profile of surface actin intensity normalized to posterior-most value along the AP axis of E8.5 mouse embryos. Anterior end of PSM is shown (vertical dashed line). Data binned in 15 μm intervals. N=11 embryos. Mean and sem are shown for all quantifications. Scale bars, 100 μm.

## Human and mouse gastruloids share the same physical mechanism of axis elongation

To determine whether the physical mechanism of axis elongation that we identified in mouse gastruloids and embryos is conserved in humans — a question that cannot be directly addressed in human embryos due to ethical and legal limitations — we turned to human gastruloids as a model system. First, we established a protocol for reproducible gastruloid formation from human induced pluripotent stem cells by combining insights from previously published methods^28,29^ (Fig. 6a,b; Methods). Similar to mouse, human gastruloids displayed volumetric growth during elongation (Fig. 6b,c) and a posterior domain that elongated without thinning (Extended Fig. 7; Supplementary Video 8). Measurements of cell divisions showed a rather uniform density along the AP axis (Fig. 6b,d), with inhibition of cell proliferation abolishing axis elongation completely (Fig. 6e,f). Oil microdroplets inserted in the tissue elongated in the perpendicular direction to the local AP axis (Fig. 6g-h), and inhibition of non-muscle myosin II did not affect elongation (Fig. 6i-j). Together, all these findings indicate that, like for mouse gastruloids, human gastruloid elongation is not governed by CE, but instead driven by the expansive forces of proliferation. Quantification of surface actin revealed a posterior actin cap with a similar decay length scale as in mouse gastruloids (Fig. 6k-l). Treatment with Cytochalasin D disrupted the actin cap and prevented axis elongation (Fig. 6m-n). Altogether, these results show conservation of the physical mechanism of axis elongation from mouse to human.

**Figure 6.**
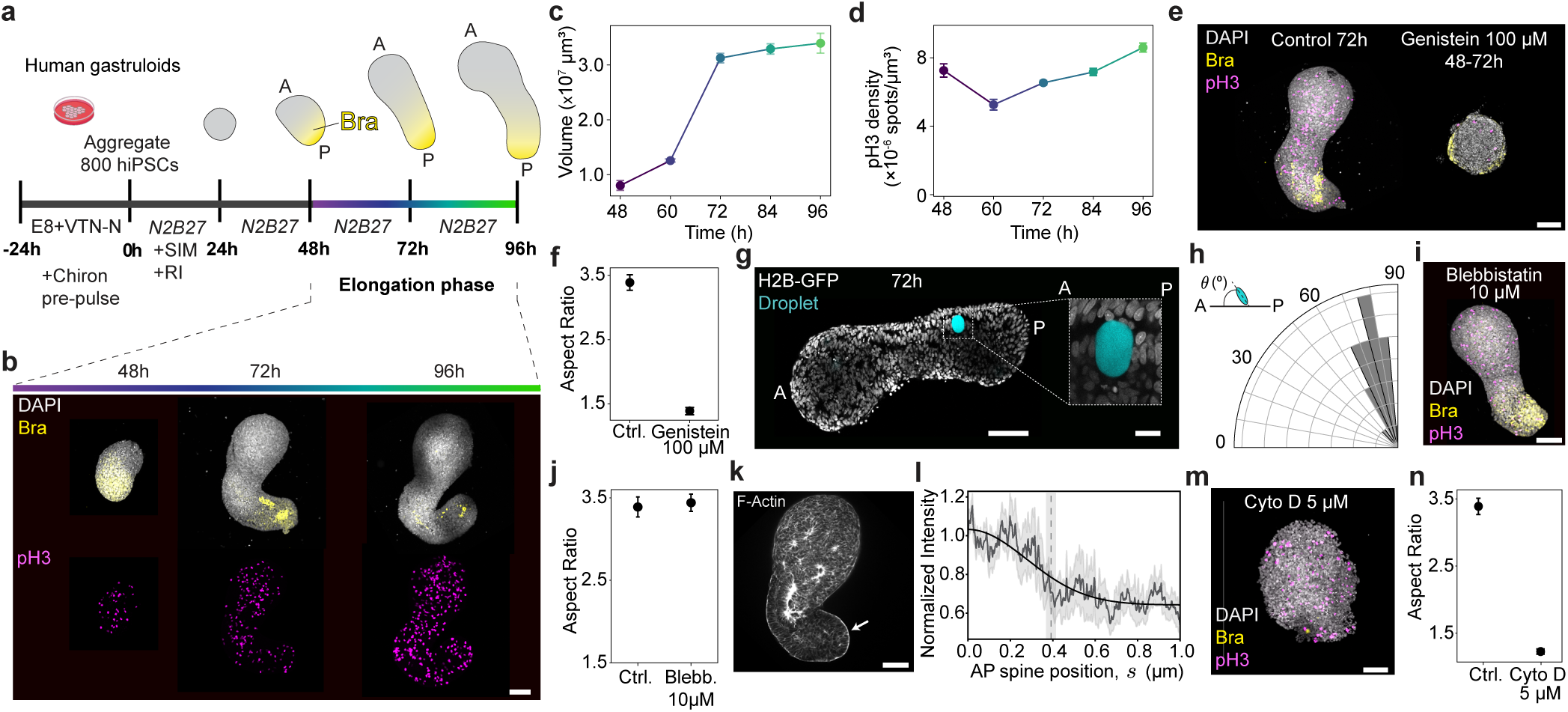
Human gastruloids elongate via the same physical mechanism as mouse gastruloids. **a**, Protocol for human gastruloid generation from hiPSCs. Bra expression (yellow) indicates posterior domain. Elongation phase occurs between 48-96h (time color coded from purple to green). **b**, Confocal images (MIPs) of fixed human gastruloids stained for DAPI and Bra (gray and yellow, respectively; top) and pH3 (magenta, bottom) at 48h, 72h and 96h after aggregation. **c**-**d**, Time evolution of gastruloid volume (**c**) and pH3 density (**d**) in human gastruloids during elongation. N= 21, 24, 20, 13, 23 for each time point, respectively. **e-f**, Confocal imaging (MIP) of fixed human gastruloids (72h) stained for DAPI, Bra and pH3 in control and Genistein 100 μM conditions (**e**), and corresponding measurements of the gastruloid aspect ratio (**f**; N = 20, 27, respectively). **g**, Confocal section of a nuclear-labelled (H2B-GFP, gray) live human gastruloid (72h) with an inserted oil droplet (cyan); closeup of the droplet shown (dashed square). **h**, Measured angle *8* between the droplet elongation axis and AP axis in human gastruloids at 72h. N= 8. **i**-**j**, Confocal image (MIP) of a human gastruloid treated with Blebbistatin 10 μM between 48-72h and stained for DAPI, Bra and pH3 (**i**) and associated quantification of gastruloid aspect ratio in control and Blebbistatin-treated conditions (**j**; N = 20, 24 respectively). **k**-**l**, Confocal section of a fixed human gastruloid stained for F-Actin at a 72h (**k**), and measured surface actin intensity along the AP axis normalized to its posterior values (**l**; mean, gray line; s.e.m, gray band; fit, black line; decay length scale indicated as vertical dashed line). N = 6. **m-n**, Confocal image (MIP) of a fixed human gastruloid treated with 5 μM Cytochalasin D between 48-72h and stained for DAPI, Bra, pH3 (**m**) and associated quantification of gastruloid aspect ratio in control and Cytochalasin D-treated conditions (**n**; N = 20, 22, respectively). Mean ± s.e.m in all quantifications. Scale bars, 100 μm, except for **g,** which has a scale of 20 μm.

## Discussion

Our results reveal a novel physical mechanism of body axis elongation in mammalian species, including humans. Cell proliferation generates the necessary forces for posterior elongation, with a posterior actin cap constraining lateral tissue expansion to ensure unidirectional posterior elongation. We observed the key features of this mechanism in mouse embryos, suggesting that this physical mechanism of axis elongation is also shared by mammalian species *in vivo*.

While convergent extension (CE) occurs in specific mouse embryonic tissues^30–35^, whether CE constitutes the mechanism underlying mammalian body axis elongation so far has remained unresolved. As opposed to other amniotes, no CE behaviors are observed during mouse primitive streak formation^36^ . Theory can explain gastruloid shapes by CE^37^ and recent work suggests a role for WNT/PCP in gastruloid elongation, but no clear signatures of CE have been observed^38^. We provide direct proof that mammalian body axis elongation can proceed independently of classical CE mechanisms: our direct force measurements show the absence of the characteristic mediolateral stresses that drive CE movements; our geometrical 3D measurements do not show any thinning during elongation; and elongation proceeds upon myosin II inhibition. Our results indicate that mouse embryos use the same mechanism as gastruloids to elongate, with inhibition of cell proliferation precluding axis elongation. This is further supported by *in vivo* observations showing that — at elongation stages beyond those modelled by gastruloids — both division rates and cell cycle duration are similar along the AP axis^39^ , suggesting constant expansion of the tailbud and PSM region. However, we cannot completely exclude a partial contribution of CE to posterior elongation *in vivo*. Likewise, given that gastruloids lack a neural tube and notochord^15,16,28^, tissues for which CE has been observed during mouse axis elongation *in vivo*^30–35^, the induction of neural tube^16^ and/or notochord^40^ formation in gastruloids might trigger CE-like behaviors.

Different vertebrate species seem to have evolved distinct physical mechanisms of elongation^3,4^. Previously reported physical mechanisms of body axis elongation in chick and fish rely on AP gradients of bulk tissue material properties or material (fluid/solid) transitions^7,8,41,42^. However, unlike fish and chick^43^, we find that in mammals — for which body axis elongation coincides with posterior volumetric growth^30,43,44^ — cell proliferation is the driving force for elongation, with cell divisions being uniform in the tissue. Since cell divisions are known to relax mechanical stresses and fluidize tissues^45–47^, it is likely that mouse and human gastruloids, as well as the mouse PSM, display a uniform bulk tissue fluidity controlled by cell division events. Even with uniform bulk tissue fluidity, surface mechanical constraints can redirect isotropic expansion into elongation, as observed in other species, such as *C. elegans* or even plants and fungi^3^. In these organisms, isotropic pressure (either hydrostatic pressure or tissue pressure generated by proliferation) is constrained laterally to guide the unidirectional elongation of the organism. Our results indicate that mammalian species use a conceptually similar solution, albeit with a unique implementation: a posterior, supracellular actin cap at the tissue surface constraining the isotropic expansion forces of uniform cell proliferation. Therefore, the control of surface tissue mechanics plays a central role in mammalian axis elongation, adding to its key role in the initial symmetry breaking process of gastruloids^48^.

Altogether, our data reveal that mammalian species, including humans, physically build their body axis via a different mechanism than other vertebrate species. More generally, these findings highlight that, across vertebrates and throughout evolution, distinct physical quantities are controlled to produce similar body plans.

## Acknowledgements

We thank the members of the Veenvliet and Campàs labs for helpful discussions, the Light Microscopy, Organoid & Stem Cell, Genome Engineering and Transgenic Core facilities of the Max Planck Institute of Molecular Cell Biology and Genetics (MPI-CBG), as well as its Biomedical Services and Technology Development Studio for their technical support. We thank Adriano Bolondi, Julia Batki, and Alexander Meissner (Max Planck Institute for Molecular Genetics, Berlin, Germany) for kindly gifting us the NLS-GFP mESC line, Shuo-Ting Yen for help with 3D printing, and the Brugués lab for sharing knowledge and materials for the design of light-sheet microwell molds. MTB is supported by a Boehringer Ingelheim Fonds PhD Fellowship. This work was supported by the Max Planck Society, a European Innovation Council (EIC) Pathfinder grant under the Horizon Europe Research and Innovation Program (Horizon-EIC-2021-PathfinderChallenges-01 101071203, SUMO), the Stiftung zur Förderung und Erforschung von Ersatz-und Ergänzungsmethoden zur Einschränkung von Tierversuchen (SET), and the Deutsche Forschungsgemeinschaft (DFG, German Research Foundation) under Germany’s Excellence Strategy – EXC 2068 – 390729961– Cluster of Excellence Physics of Life of TU Dresden.

## Author Contributions Statement

JVV and OC conceptualized the project. MTB, JVV and OC designed research; MTB performed all gastruloid and mouse embryo experiments; RGS developed data analysis codes; MTB, RGS and JRS analyzed data; AB and MTB performed and analyzed laser ablation experiments; CF set up injection system for gastruloids and performed microdroplet calibrations; PP designed molds for lightsheet imaging; MTB set up the human gastruloid protocol, with initial help from DC; HLvdW and EMS provided custom-made surfactants. MTB, JVV and OC wrote the paper, with input from RGS, AB and PP; JVV and OC supervised the project.

## Competing Interests Statement

The authors declare that they have no competing interests of any kind.

## Extended Figure Legends

**Extended Figure 1.**
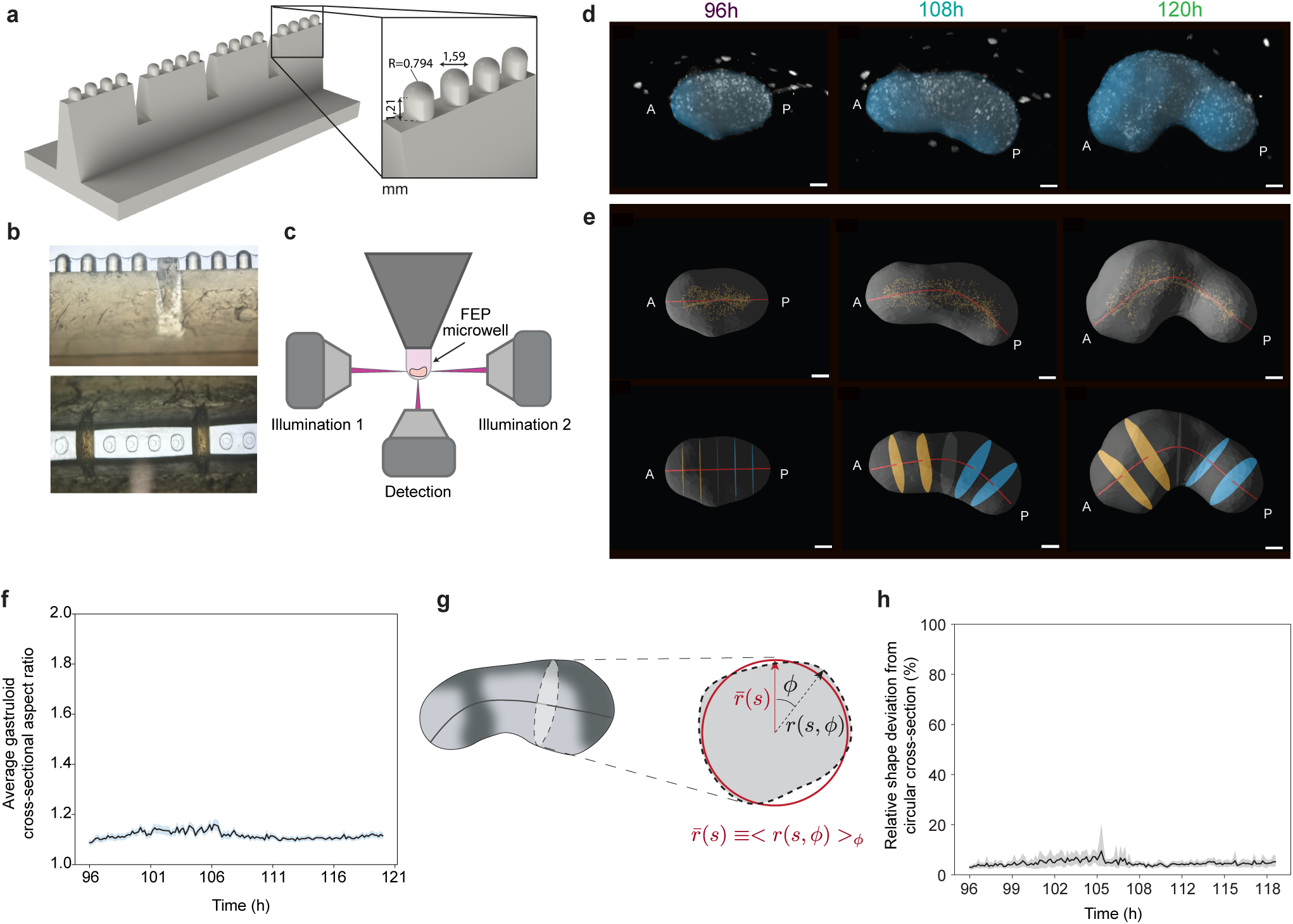
Light-sheet live imaging setup for mouse gastruloids and geometrical characterization of gastruloids. **a**-**c**, Microwell mold design for simultaneous live imaging of up to 16 gastruloids with a Viventis light-sheet microscope. Molds were 3D printed and used for replicating microwells onto FEP cuvettes. Mold dimensions are shown in mm (**a**). Example mold microwells (**b**) and light-sheet setup with dual illumination (**c**) are shown. **d**-**e**, Representative live mouse gastruloid 3D images at 96h, 108h and 120h, showing the smoothed surface volumes (blue, **d**), spine detection (red, **e**) and orthogonal slices along the spine for diameter measurements at anterior (orange, **e**) and posterior (blue, **e**) regions over time. Scale bars, 100 μm. **f**, temporal evolution of the average cross-sectional aspect ratio of gastruloid slices orthogonal to the AP axis in live mouse gastruloids, showing that throughout the AP axis the gastruloid diameter maintains an approximately constant aspect ratio close to 1, indicating axisymmetry during elongation. **g**-**h**, Measurements of the deviation in shape between orthogonal gastruloid slices along the AP axis and a circle, with the shape of the gastruloid orthogonal slice at a position *s* along the spine given by *r*(*s*, *ϕ*). The relative error of the shape of orthogonal AP slices from circular cross-sections was between 5-10% throughout the 24h time window of elongation. N=12 live imaged mouse gastruloids for 24h for all. Mean lines with s.e.m. shadows are shown.

**Extended Figure 2.**
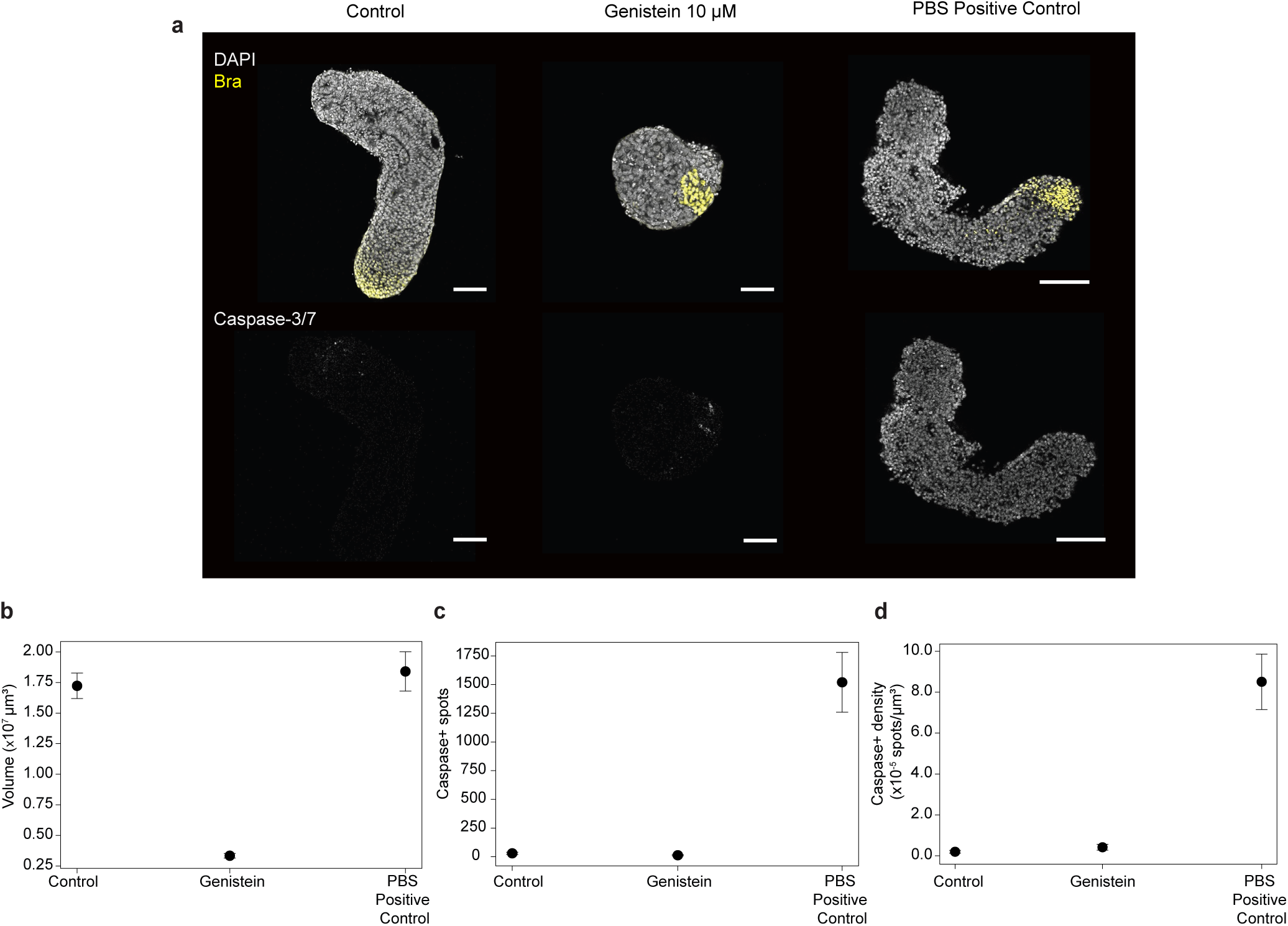
Genistein treatments in gastruloids block proliferation without inducing cell death. **a**, Representative stainings for DAPI, Bra and cleaved Caspase-3/7 (marker of apoptosis) of 120h mouse gastruloids in control, 10 μM Genistein and PBS incubation (positive control for cell death) conditions. Cleaved Caspase-3/7 positivity appears prominent in samples incubated with PBS for 1h instead of medium, but not in control or Genistein conditions, showing that Genistein treatment does not cause cell death. In conditions where cell death is induced (PBS positive control), tissue structure and nuclear morphologies appear perturbed, resembling apoptotic cells, but not in negative controls or in Genistein treatments. Single slices are shown for the different conditions. Scale bars, 100 μm. **b**-**d**, Quantifications of gastruloid volumes (**b**), number of Caspase+ nuclei (spots; **c**) and Caspase+ nuclear density (**d**) in control, Genistein, and PBS Positive control conditions. Mean and s.e.m error bars are shown for all quantifications. N = 16, 15, 11 for negative control, Genistein and positive control conditions, respectively.

**Extended Figure 3.**
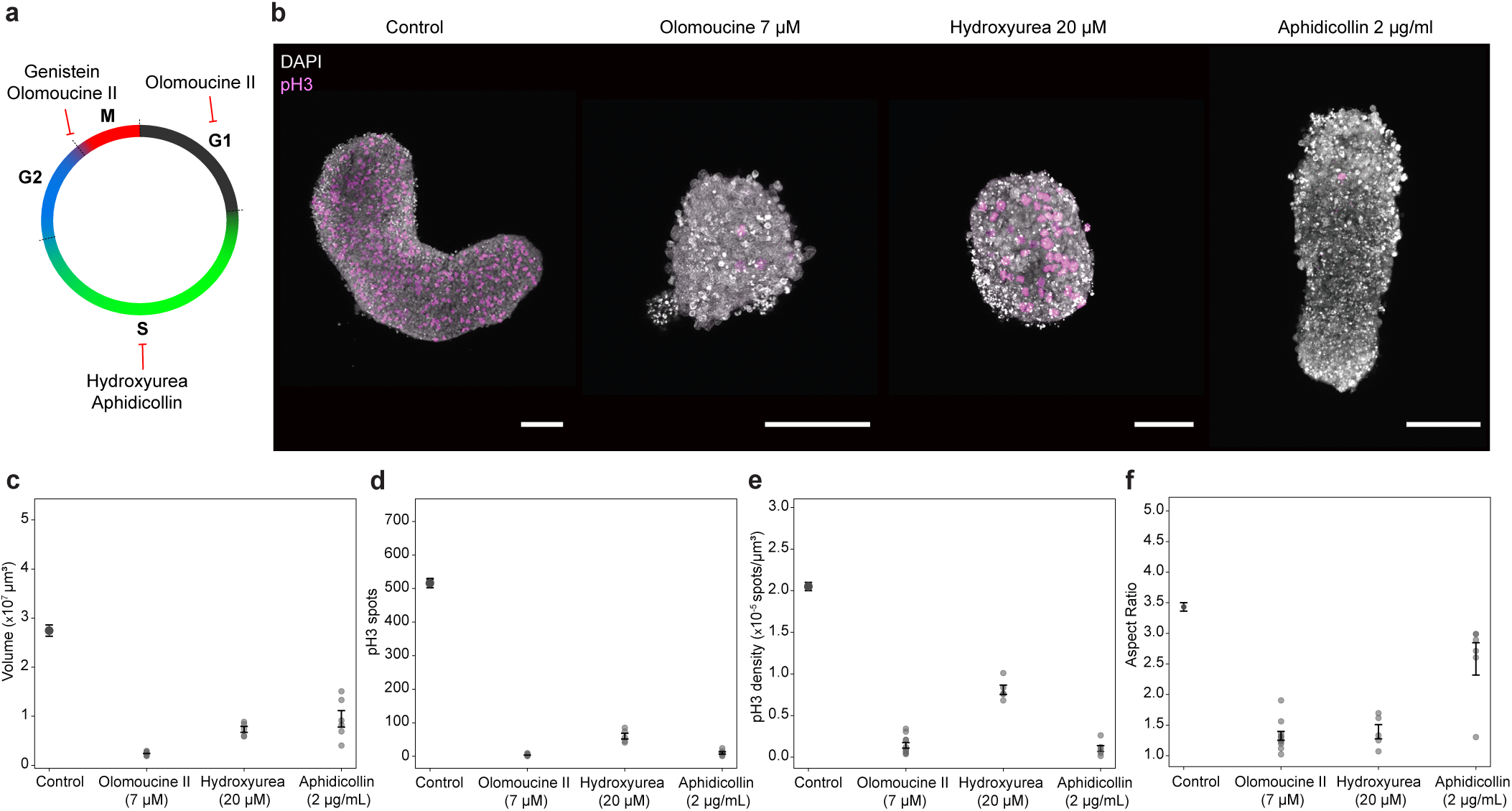
Blocking proliferation with multiple cell cycle inhibitors shows similar effects on volumetric growth and elongation in mouse gastruloids. **a**, Schematic showing the effect of each inhibitor used for blocking proliferation on specific phases of the cell cycle. Genistein is known to block the entry into mitosis (M phase), while Hydroxyurea and Aphidicollin are known to block DNA synthesis and progression from S phase. Olomoucine II has been reported to block both the entry into mitosis as well as the progression into the S phase. **b**, Stainings for DAPI and pH3 of 120h gastruloids in control, Olomoucine II 7 μM, Hydroxyurea 20 μM and Aphidicollin 2 μg/ml conditions. A clear reduction in the elongation levels and pH3 positivity is observed in gastruloids treated with the cell cycle inhibitors between 96h-120h compared to control gastruloids at 120h. Max projections shown. Scale bars, 100 μm. **c**-**f**, Quantifications of total volumes (**c**), pH3+ nuclei (spots; **d**), pH3 densities (**e**) and aspect ratio (**f**) in control and in Olomoucine II, Hydroxyurea and Aphidicollin-treated gastruloids at 120h. A reduction in all parameters related to mouse gastruloid elongation (proliferation, volumetric growth and aspect ratio) was observed with all cell cycle inhibitor treatments. Mean and s.e.m error bars are shown for all quantifications. N = 13, 11, 5, 6 for Control, Olomoucine II, Hydroxyurea and Aphidicollin, respectively.

**Extended Figure 4.**
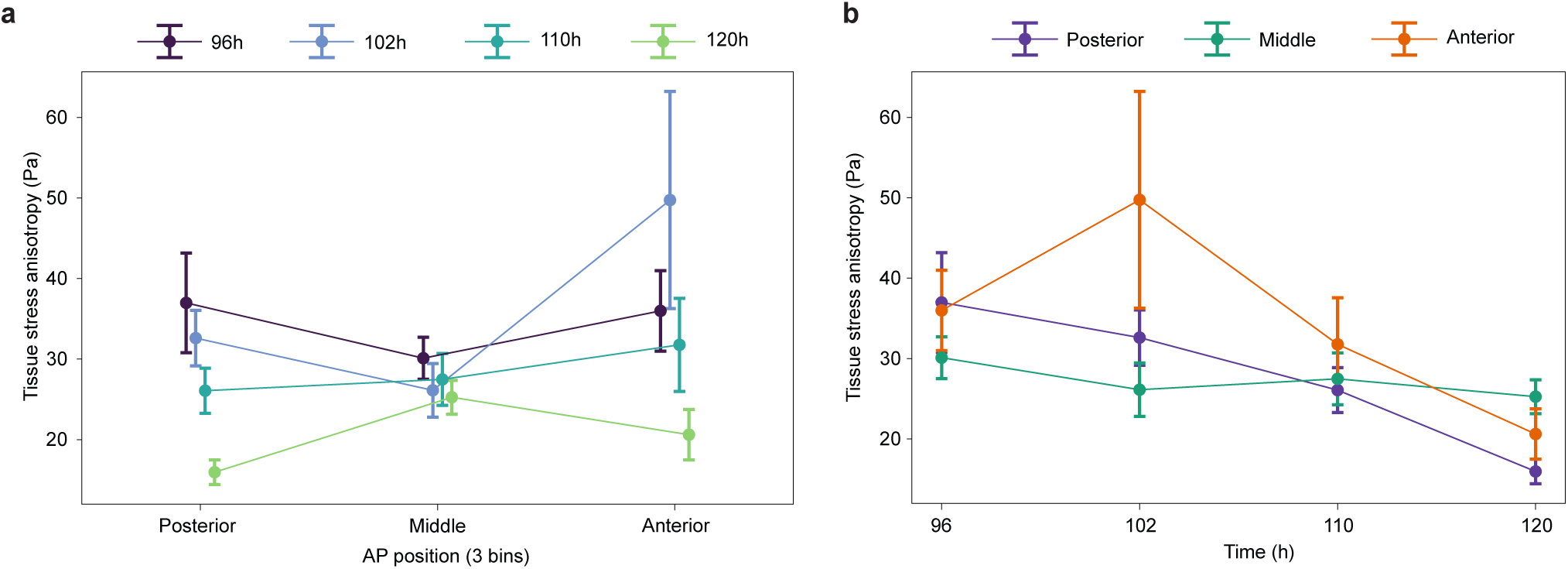
Spatiotemporal variations in tissue-scale stress anisotropy during mouse gastruloid elongation. **a-b**, Measurements of tissue stresses from droplets injected and imaged along the AP axis at each time point of elongation, allowing for the spatiotemporal mapping of tissue stresses throughout elongation. After spine detection from largeview images of gastruloids and droplet centroid and main axis computation from napari-STRESS analysis of droplets, each droplet is assigned an AP axis position by dividing the AP axis in three equal bins (Posterior, Middle and Anterior regions). Tissue stresses are measured for each region and for each time point. **a**, Tissue-scale stress anisotropy along the AP axis: Posterior (purple), Middle (green) and Anterior (Orange), for the different timepoints during elongation. **b**, Temporal evolution of tissue-scale stress anisotropy in each region along the AP axis. Mean and s.e.m error bars are shown for all quantifications. For Anterior, Middle and Posterior regions, respectively, N = 26, 34, 22 for 96h gastruloids; N = 26, 17, 6 for 102h Gastruloids; N = 5, 29, 8 for 110h Gastruloids, and N = 10, 3, 6 for 120h Gastruloids.

**Extended Figure 5.**
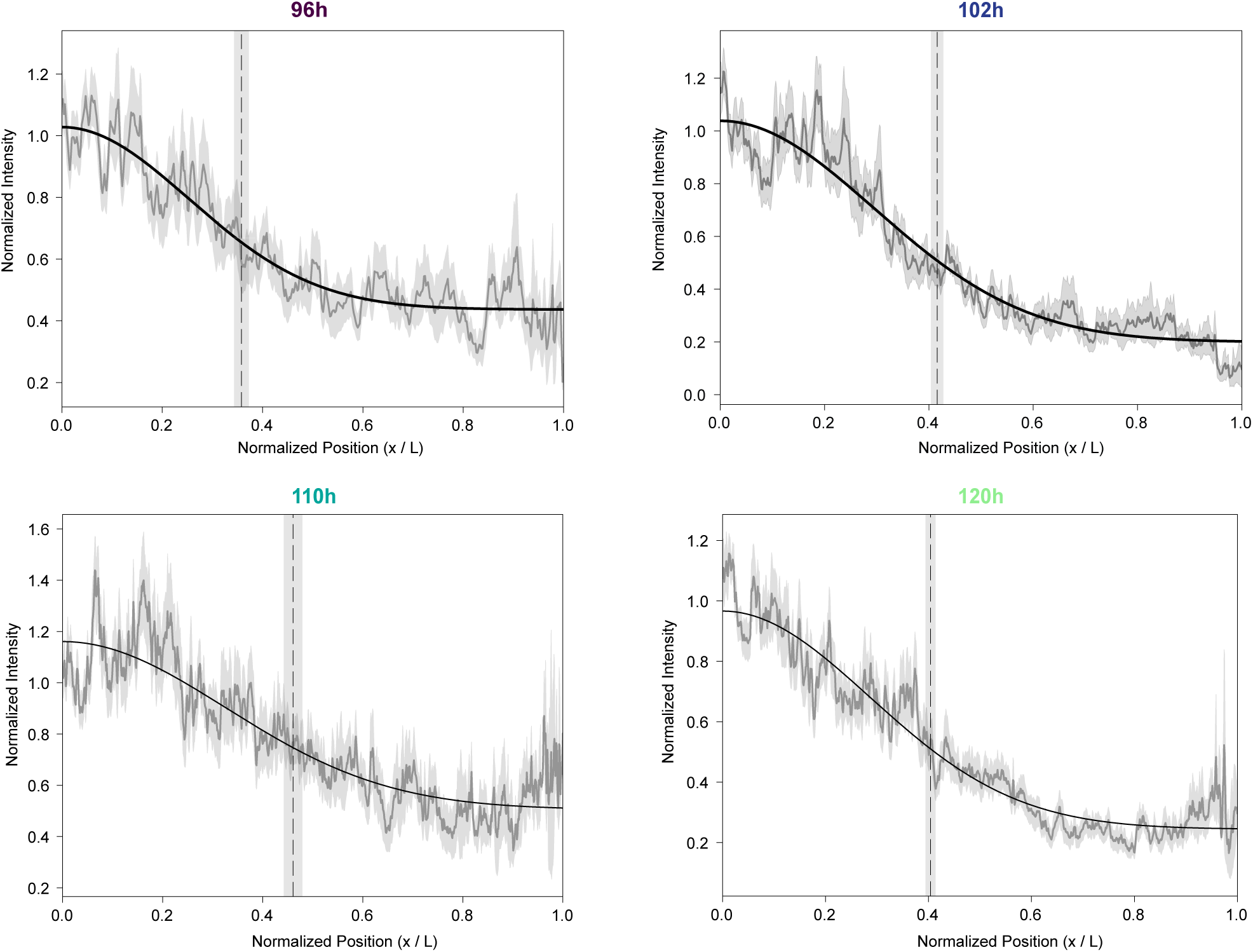
Time evolution of the actin cap during elongation of mouse gastruloids. Normalized surface F-Actin intensity decay along the normalized AP axis (x/L) length for each mouse gastruloid time point (96h, 102h, 110h, 120h, indicated with purple-to-green color gradient as in main figures), corresponding to the surface actin intensity decay measurements in Fig. 4d, e. Actin decay fits are indicated with gray lines, and actin decay lengthscale (α) for each time point are indicated with a dashed vertical line and a shadowed error bar from fit residuals. Mean lines with s.e.m. shadows are plotted as in Fig.4. N = 11, 11, 12, 12 for each time point, respectively, as in Fig. 4.

**Extended Figure 6.**
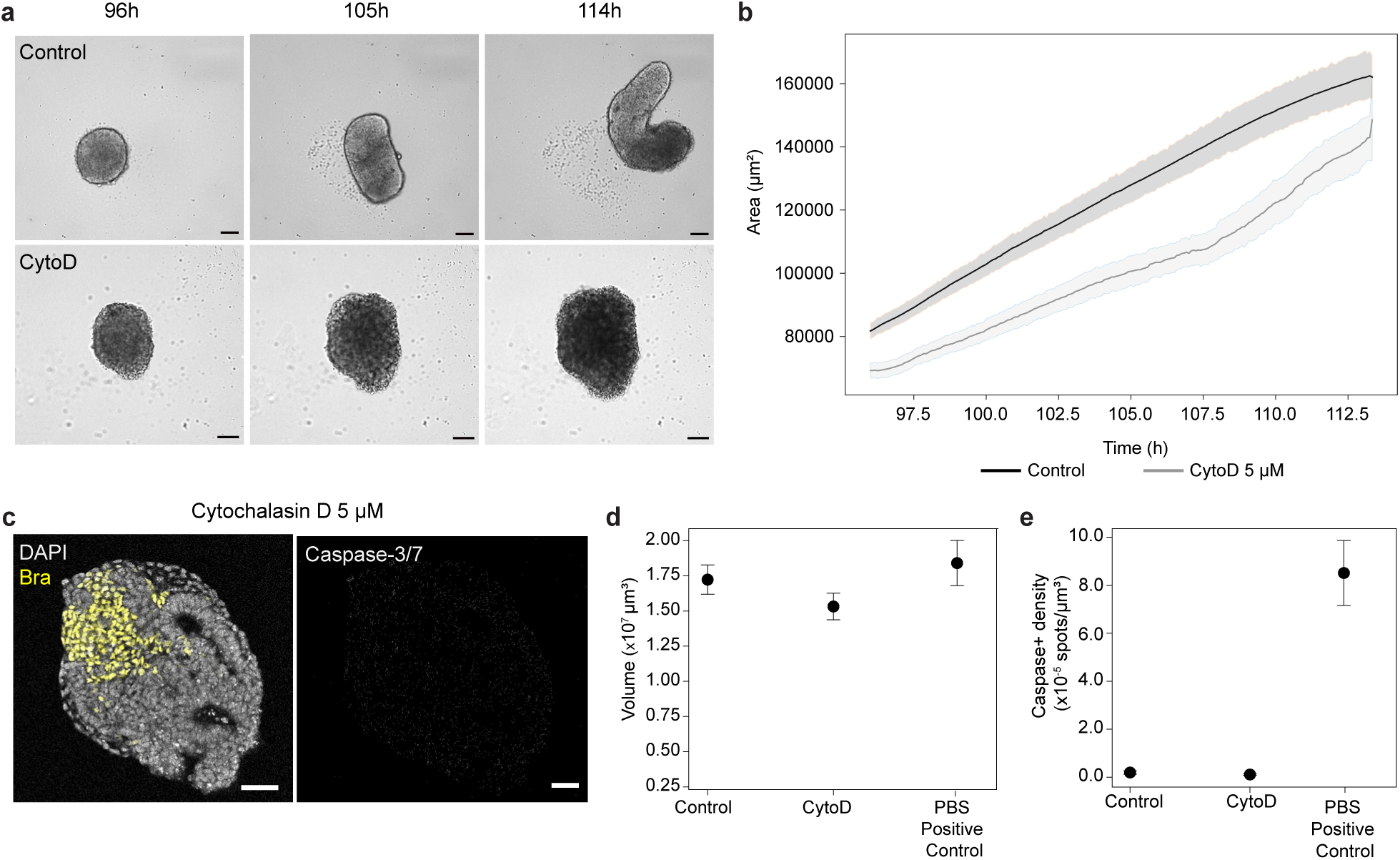
Characterization of Cytochalasin D effects on elongation in mouse gastruloids. **a**, Brightfield live images of mouse gastruloids at 96h, 105h and 114h in control and in 5 μM Cytochalasin D conditions from 2D live imaging of gastruloid development during the elongation phase. Control gastruloids follow tube-like elongation dynamics, while Cyto D-treated gastruloids expand isotropically. Scale bars, 100 μm. **b**, Measurements of gastruloid 2D projected area over time from brightfield live imaging during the elongation phase in control (black) and Cyto D-treated gastruloids (gray), showing an increase in Area (and therefore of corresponding Volume in 3D) for both conditions. Mean lines with s.e.m bands are shown. N = 32 for each condition. **c**, Staining for DAPI, Bra and cleaved Caspase-3/7 (apoptosis marker) of a 120h mouse gastruloid treated with 5 μM Cytochalasin D from 96-120h. Scale bars, 100 μm. **d**-**e**, Quantifications of gastruloids total volumes (**d**) and Caspase-3/7+ nuclei densities (**e**) in negative control, Cyto D-treated, and PBS-treated positive control (1h) 120h Gastruloids. While all conditions present volumetric growth, and similar levels of volume by 120h of culture, only the positive control (incubation in PBS for 1h) presents high apoptosis from Caspase-3/7+ nuclei density. Therefore, Cyto D treatment does not seem to induce apoptosis in mouse gastruloids. Mean and s.e.m error bars are shown for all quantifications. N = 16, 12, 11 for negative control, Cyto D and positive control conditions, respectively.

**Extended Figure 7.**
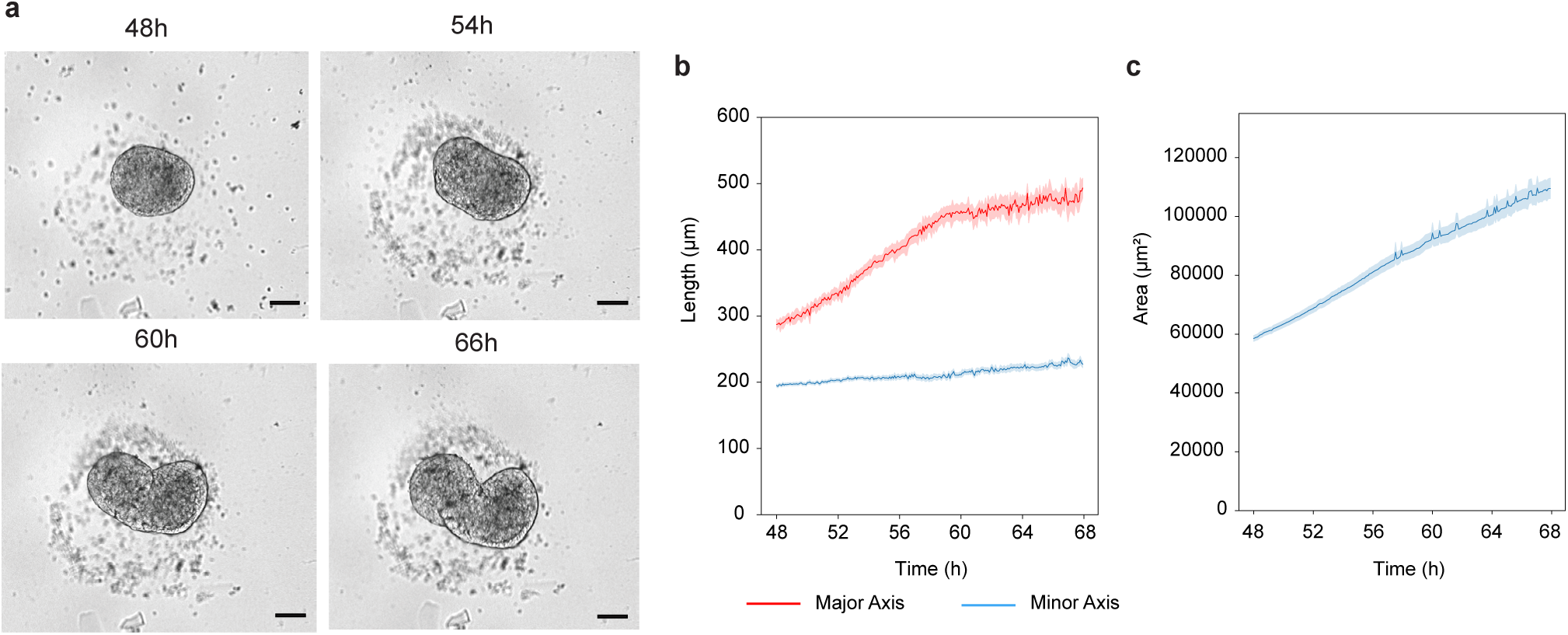
Axial elongation dynamics of human gastruloids from brightfield imaging. **a**, Representative brightfield images of a 2D live imaging experiment of human gastruloid axial elongation between 48-72h. Scale bars, 100 μm. **b**-**c**, Measurements of major axis (red; **b**) and minor axis (blue; **b**), as well as area changes (**c**) over time for live imaged human gastruloids in 2D during the elongation phase. While total structure length keeps increasing during elongation, as does the total gastruloids area, the minor axis seems to be maintained constant throughout elongation, similarly as in mouse gastruloids, indicating a lack of tissue thinning and convergent extension behaviors in the elongation of human gastruloids. Mean lines with s.e.m error bars are plotted. N = 29 live imaged human gastruloids.

## Methods

### Mouse husbandry, embryo collection and tailbud explant culture

Mouse embryos were collected from adult female C57BL/6JOlaHsd mice in the animal facility of the Max Planck Institute of Molecular Cell Biology and Genetics (MPI-CBG), Dresden, Germany, in accordance with the German Animal Welfare Legislation after approval by the federal state authority Landesdirektion Sachsen (licenses: TVT07/2018 and TVT08/2023). All procedures were overseen by the Institutional Animal Welfare Officer and the institutional Animal Welfare Body. All animals used for this study were kept in standardized pathogen-free conditions at the Biomedical Services Facility (BMS) of MPI-CBG with free access to food and water in a 12 h/12 h light/dark cycle. Adult females were time-mated and checked for vaginal plugs one day after mating. Plugged females were humanely killed for embryo collection by cervical dislocation at E8.5. Mouse embryos were collected and dissected from uterine horns in PBS under a stereoscope, removing extraembryonic membranes using forceps. Mouse embryos were fixed after dissection, by collecting them in 8-well dishes (Ibidi 80860), washing three times in PBS, fixing in 4% PFA (Sigma, 158127) for 30 min, and washing three times more in PBS. For tail explant culture experiments, E8.5 mouse embryo tails were cut at somite 4 using forceps, and explants were collected in 8-well dishes in Ndiff227 medium (Takara, Y40002) with 1X penicillin (5000 U/ml)/ streptomycin (5000 μg/ml) (Sigma, A8674) and placed in an incubator at 37°C and 5% CO_2_.

### Mouse embryonic stem cell culture

Mouse embryonic stem cells (mESC) with F1G4 genetic background^49^ were cultured in 6-well tissue culture plates (Nunc™, TFS 140675) coated with 0.1% Gelatin (Sigma G1393) and with a mitotically inactive mouse embryonic fibroblast layer, in standard ES+LIF medium. WT F1G4 mESCs, or F1G4 carrying a NLS-GFP transgene inserted in the *Rosa26* locus^20^ (kindly provided by the Meissner lab) or F1G4 carrying a Lifeact-mScarlet transgene inserted in the *Rosa26* locus (see below) were used. ES + LIF medium was prepared as follows: for 500 ml of medium, mix 400 ml Knockout Dulbecco’s Modified Eagle’s Medium (DMEM) (4500 mg/ml glucose, w/o sodium pyruvate) (Gibco, 11965092) with 75 ml ES cell tested FCS, 5 ml 100x L-glutamine (200 nM) (Lonza, BE17-605E), 5 mL 100x penicillin (5000 U/ml)/ streptomycin (5000 μg/ml) (Sigma, A8674), 5 ml 100x non-essential amino acids (Gibco, 11140-35), 1 mL 500x β-mercaptoethanol (5mM, Invitrogen, 21985023) and 5 mL 100x nucleosides (Chemicon, ES-008-D). Medium was aliquoted and stored at -20°C until use, when medium was thawed and supplemented with 1:10000 of Murine Leukemia Inhibitory Factor (LIF) (Sigma-Aldrich, ESG1107). mESCs were cultured at a starting density of 80,000-100,000 cells/well after thawing and passaging, with daily media changes and passaging every other day using Trypsin-EDTA (TFS, 25300054). Passaging was done at least two times before starting a gastruloid generation. Cultures were routinely confirmed to be negative for mycoplasma by PCR of the medium supernatant after three days without medium change.

### Generation of Lifeact-mScarlet mESCs

Lifeact-mScarlet was inserted as a transgene by CRISPR/Cas9 into the mouse Rosa26 locus. mRosa26 guideRNA (AGTCTTCTGGGCAGGCTTAA) was ordered as crRNA from IDT (Integrated DNA Technologies). crRNAs were duplexed with tracrRNA (IDT Cat.no. 1072534) by mixing equimolar amounts. The targeting construct containing a neomycin resistance selection cassette and CAG promotor driven Lifeact-mScarlet CDS flanked by 500 bp homology arms was generated through Gene Synthesis by Genscript and subsequenct Type II restriction enzyme cloning. 2x 10^5 F1G4 mES cells were co-transfected with RNPs (44 pmol cr:tracr duplexes assembled with 37 pmol Cas9, IDT Cat.no. 1081061) and 500 ng of Rosa26 targeting construct using the NEON transfection system (Thermo fisher scientific). Single-cell clones were picked after selection with G418 (200 µg/ml) for 3 days and cultured in 96-well plates for 4 more days in ES-cell media (Knock-out DMEM, 20% (v/v) FCS, 2-4 mM L-Glutamin, 1% (v/v) penicillin/streptomycin, 0.1 mM 2-Mercaptoethanol, 1x non-essential amino acids, 1000 U/ml LIF) at 37 °C in an incubator with 20% O2 and 5% CO2. Genomic DNA from single-cell clones was extracted using the QuickExtract DNA extraction kit (Epicentre) following the manufacturer’s instructions. Lifeact-mScarlet single-cell clones were identified by PCR using Phusion Flash High-Fidelity PCR Master Mix (ThermoFisher) and verified by Sanger Sequencing (primer sequences see table below). Verified clones were thawed and expanded for further analysis.

### Generation of mouse gastruloids

Gastruloid generation was performed as previously described^16,50^ . In brief, mESCs were feeder-freed by plating in 0.1% gelatin 6-well plates after passaging, and sequentially incubating at 37°C for 25, 20 and 15 min. Feeder-freed mESCs were then collected by centrifuging 5’ at 300g, then washed with 5 ml of pre-warmed PBS with MgCl_2_ and CaCl_2_ (PBS+\+) (Sigma, D8662-500ml), pelleted (5’ 300g) and washed again with 5 ml pre-incubated Ndiff227 medium (Takara). After centrifuging again (5’ 300g), mESCs were resuspended in 200-300 μl of pre-warmed Ndiff227 and counted with an automated cell counter (Invitrogen™ Countess™ 3 FL Automated Cell Counter). mESCs were then diluted in 5 ml of NDiff227 to a final concentration of 8,571 cells/ml. We then plated 35 μl/well in an ultra-low attachment U-bottom 96-well plate (Corning 7007), corresponding to 300 cells/well, using a multi-channel pipette, and placed plates in an incubator set at 37°C and 5% CO_2_ for 48h. After 48h, 150μl of NDiff227 medium supplemented with 3 μM CHIR (Tocris, 4423) was added to each well. Subsequently, Ndiff227 medium was refreshed every 24h by removing 150μl from each well and adding 150μl of fresh Ndiff227 pre-incubated medium.

### Human induced pluripotent stem cell culture

A human induced pluripotent stem cell (hiPSC) line with WTC-11 genetic background carrying a mEGP-H2B type 1-J tag (UCSFi001-A-28 (RRID:CVCL_UD17)) was acquired from Cornell Institute for Medical Research (CIMR) via the Organoid and Stem Cell Facility of MPI-CBG. hiPSCs were routinely cultured in 6-well plates (Nunc™, TFS 140675) coated with 5 μg/ml of rhVitronectin-N (VTN-N) (TFS, A14700) with Essential 8 (E8) medium (TFS, A1517001). VTN-N coating was performed by resuspending stock solution in PBS at a 1:100 dilution, adding 1 ml of solution to each well of a 6-well plate and incubating at room temperature for 1h. After 1h, protein solution was aspirated and E8 medium was added to each well. hiPSCs were thawed in E8 medium supplemented with 10 μM Rock inhibitor (Y-27632, STEMCELL™ Technologies, 72308) for 24h, after which daily E8 media changes were performed without Rock inhibitor. Passaging was done when 80% confluency was reached, using ReLeSR™ (STEMCELL™ Technologies, 100-0483), by incubating with the reagent for 1 min at room temperature, then aspirating it and incubating at 37°C for 5 min. After that, cells were resuspended in 1 ml of E8 medium into clumps without extensive pipetting, and cell clumps were passaged at a 1:8-1:10 ratio into pre-coated wells. At least two passages were performed before starting a human gastruloid generation. Cultures were confirmed to be negative for mycoplasma by PCR of the medium supernatant after three days without medium change.

### Generation of human gastruloids

Human gastruloid generations were performed with a protocol adapted from previous studies^28,29^ . hiPSC cultures used for a generation were pre-pulsed with E8 medium supplemented with 3 μM CHIR (Tocris #4423) for 24h (when confluency reached about 60%). After 24h, hiPSCs were washed with PBS, dissociated into single cells with Trypsin-EDTA (TFS, 25300054), and resuspended in E8 medium supplemented with 10 μM Rock inhibitor (Y-27632). Cells were centrifuged 5 min at 300g, washed once with 5 ml of homemade N2B27 medium supplemented with 10 μM Y-27632, centrifuged 5 min at 300g, and finally resuspended in 500 μl of N2B27 with 10 μM Y-27632 and Somite Induction Factors (SIM^29^ ). Cells were counted with an automated cell counter (Invitrogen™ Countess™ 3 FL Automated Cell Counter), and a cell suspension was made in 6 ml of N2B27 with 10 μM Y-27632 and SIM factors to reach a final density of 16,000 cells/ml. 50 μl of this cell suspension was transferred into each well of an ultra-low adherence U-bottom 96-well microplate (Nunclon Sphera, TFS #174925), corresponding to 800 cell aggregates/well. Plates were then centrifuged 3 min at 300g and kept in an incubator set at 37°C and 5% CO_2_. After 24h, 150μl of pre-incubated N2B27 medium were added to each well, and every 24h thereafter medium was refreshed by removing 150μl from each well and adding 150μl of pre-incubated N2B27, until 96h after aggregation. Homemade N2B27 medium was prepared as follows: for 200 ml of medium, mix 94.5 ml of DMEM/F12 (Gibco, 21331020) and 94.5 ml of Neurobasal (Gibco, 21203049), 2ml of 100X penicillin (5000 U/ml)/ streptomycin (5000 μg/ml) (Sigma, A8674), 2 ml of 200mM Glutamax (TFS, 35050061), 2 ml of 100X non-essential amino acids (Gibco, 11140-35), 2 ml of 100mM sodium pyruvate (TFS, 11360070), 1 ml of 100X N2 supplement (R&D, AR009), 2 ml of 50X B27 supplement with Vitamin A (Gibco, 17504-044) and 0.8 ml of beta-mercaptoethanol (Gibco, 21985-023). SIM factors were added to N2B27 at the following concentrations: 10 μM SB431542 (Sigma, S4317), 5 μM CHIR (Tocris, #4423), 2 μM DMH1 (Sigma, D8946) and 20 ng/ml bFGF (Peprotech, AF-100-18B). All human gastruloid experiments followed the Guidelines for Stem Cell Research and Clinical Translation published by the International Society for Stem Cell Research. Approval for the generation of human gastruloids from hiPSCs was granted by the Ethics Council of the Max Planck Society on June 21, 2023 (application ID: 2022_17).

### Whole mount immunofluorescent staining of gastruloids and mouse embryos

Fixation and staining of mouse gastruloids, human gastruloids and mouse embryos was performed as previously described^16^ . Samples were transferred into 8-well Ibidi dishes (80860) using p200 wide-bore pipette tips, washed three time with PBS, fixed with 4% PFA for 30 min, and washed three more times with PBS. Structures were permeabilized with PBST (PBS with 0.05% v/v Triton-X) for one hour, and then incubated with blocking buffer (PBST with 10% FBS) overnight at 4°C. After blocking, primary antibodies were added in blocking buffer and incubated for 3 to 7 days at 4°C. Primary antibodies used were rabbit anti-Brachyury (1:250, CST 81694S) and rat anti-phospho-Histone3 (1:300, Sigma H9908). After primary antibody incubation, structures were washed three times with PBST and three times with blocking buffer, and then blocked overnight in blocking buffer at 4°C. Subsequently, secondary antibodies were added in blocking buffer at a 1:500 dilution and incubated overnight at 4°C, together with DAPI-405 (1:2000, Merck Roche 10236276001) and Phalloidin-488 (1:400, TFS A12379). Secondary antibodies used were donkey anti-rabbit-568 (TFS A10042), donkey anti-goat-405 (TFS A48259) and donkey anti-rat-647 (TFS A48272). For cleaved Caspase 3/7 stainings, the Cell event caspase kit (TFS C10723) was used, incubating live samples for 1h with a 1:1000 dilution of the conjugated antibody, and then proceeding with the fixation and staining steps explained above. After that, structures were washed three times with blocking buffer and three times with PBS, followed by a post-staining fixation in 4% PFA for 30 min. Finally, structures were washed two times in 0.02M Phosphate Buffer (PB) and mounted with a small drop of 1% low-melting point agarose in 0.02M PB to stabilize structures for imaging. After agarose solidified, structures were cleared by adding RIMS, consisting of 133% w/v Histodenz (Sigma D2138) in 0.02 M PB, and incubating overnight at 4°C before imaging.

### Imaging of gastruloids and mouse embryos

Live imaging of gastruloids was performed using a Viventis LS1 light sheet microscope. Custom-made 3D printed molds were fabricated to generate microwells for imaging of free-floating gastruloids in the Viventis FEP cuvette (Extended Fig. 1a). In short, molds of the size of Viventis cuvettes with 16 U-shaped microwells were designed using AutoDesk Fusion and 3D printed using a stereolithography resin 3D printer (Formlabs, Form3B+). 3D printing was done with Durable v2 resin (Formlabs) with 0.050 mm resolution steps. Molds were used to extrude microwells onto the Viventis FEP cuvettes by mechanically pressing the molds onto the FEP foil. Microwell cuvettes were washed with 70% EtOH, water, and 70% EtOH, and then exposed to UV for 30 min under a tissue culture hood. After sterilization, the molds were rinsed with anti-adherence solution (STEMCELL™ Technologies, 07010) for at least 30 min, then washed three times with NDiff227 medium, and placed in an incubator for pre-warming. Gastruloids were transferred from 96-well plates after 94h of aggregation into the Viventis cuvettes using p200 wide-bore pipette tips, and positioned for imaging into the microwells under a stereoscope in a flow hood. Gastruloids were imaged in the Viventis LS1 chamber at 37°C and 5% CO_2_ incubation, using the 488 nm laser line set at the minimal power possible for imaging of structures (20-30%), with a light sheet thickness of approximately 2.2 μm (FWHM), and a Nikon Apo 25x (1.1 NA) water immersion objective. An OD2 filter (making the measured effective output power of mW) and a GFP 525/50 BP filter, as well as an Andor Zyla sCMOS camera (VSC-12371) were used for imaging. Acquisition of images was done with 2048×048 pixels/frame (at full camera range, 0.75X zoom) and with 1 μm z-steps, imaging every 10 min for 24h.

Imaging of fixed, stained and cleared gastruloids and embryos was performed using an inverted Laser Scanning Confocal Microscope (Zeiss LSM 980) using a 20X/0.8 Apochromat (Zeiss) or a 40X/1.2 multi-immersion Aprochromat (Zeiss) objective set for Glycerin immersion (matching the refractive index of RIMS). Frames of 1024×1024 pixels and z-steps of 0.8-1 μm for the 20X imaging, and of 0.3-0.4 μm for the 40X imaging were acquired for volumetric imaging of structures (with pinhole set to 1AU), using the corresponding filters for the 405, 488, 568 and 647 nm laser lines. For Brightfield imaging of elongating gastruloids in 96-well plates and of cultured mouse embryo tailbud explants in 8-well Ibidi dishes (808060), we used an inverted Zeiss LSM980 with 37°C and 5% CO_2_ incubation. Samples were illuminated with a 488 nm laser using a 5X/0.25 Apochromat (Zeiss) objective, and the transmitted light was collected with a Non-Descanned Detector (NDD). Minimal laser power (0.3%) and minimal Gain (350 V) were used in order not to saturate the NDD. Multi-position imaging in this mode (up to 96 positions) was set for one single z-plane plane focused in the middle of each structure, acquiring frames of 1024×1024 pixels every 5 min for 24h during the elongation period (for gastruloids) or every 10 min for 4-6h for mouse embryo tailbud explants.

### Analysis of 4D gastruloid geometry

Gastruloid 3D morphodynamics during the elongation window were quantified using a novel computational framework, packaged as a modular Python toolkit (SpinePy^20^ ). Briefly, images were pre-processed by rescaling to isotropic voxel sizes of 2 μm, noise removal with a median filter of 8 μm and binarization with Otsu’s thresholding. 3D segmentation was done by binary dilation and binary opening operations, and a surface mesh was obtained with the marching cubes algorithm (sci-kit image). Surfaces were smoothed with a moving least squares approach (radius = 15, factor 0.1). Gastruloid surfaces were used to compute total volumes of structures and live volumetric changes. Surface contours were used to generate spines with a non-linear PCA approach, which defined the main anterior-posterior (AP) axis of structures and their live dynamics^20^ . Anterior and posterior regions were identified by manual annotation at the last time point of imaging, and retrospectively propagated to the first time points. Surfaces and spines were used to compute five equidistant planes along the AP axis perpendicular to the spine that define the diameters of gastruloids at anterior, middle and posterior regions. Two posterior and two anterior planes were used to compute the average posterior and anterior live diameter changes. For domain-specific dynamics of shape changes, a sphere was fitted onto the anterior slice vertices, with the centroid lying on the extracted A-P axis (spine). The posterior domain was retrieved by subtracting the diameter of the fit sphere from the anterior axis region, using the remaining axis length as posterior. To extract the anterior volumetric dynamics, we tracked the volume of the fit sphere and defined the remaining gastruloid volume as the posterior volume.

Gastruloid axisymmetry was determined by two measures. First, the coordinates of the surface intersections of the orthogonal planes were used to fit ellipsoids. The major and minor axes of these fit ellipsoids were then used to determine the cross-sectional aspect ratio. To determine the normalised errors for the measurement of radial profiles we determined the error as a function of the rotational angle *ϕ* in the orthogonal plane to the axis as 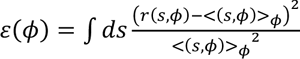 and defined the total relative shape deviation as < *ε*(*ϕ*) >*ϕ*.

### Analysis of gastruloid geometry from 2D images

To analyze 2D brightfield data, we segmented gastruloids using a custom, modified version of the ‘segment anything’ model^51^, which we fine-tuned using micro SAM^52^. Once labels were generated, we either used scikit-image to measure morphological features for the comparison of areas in mouse gastruloid cytoD perturbation experiments or a gastruloid specific framework^18^ to extract the major and minor axes in human gastruloids.

### Analysis of cell proliferation and morphometry in fixed samples

Volumetric imaging of gastruloids fixed, stained and cleared was used for measurements of total structure volumes, number of pH3+ nuclei (detected spots), density of pH3+ nuclei (spots/volume), major axis lengths, minor axis lengths and aspect ratios. Images acquired with a 20X air objective were first corrected for the refractive index (RI) mismatch between the air phase (RI = 1) and the RIMS clearing solution in which samples were embedded (RI=1.46), by converting voxel sizes using the following formula: *z_corrected* = *z*-step * (RI _RIMS_ / RI _AIR_). After this correction, 3D images were segmented using Imaris software (Oxford Instruments Imaris v10.2) based on the DAPI signal using a 10 μm surface detail with the Surfaces function, and total structure volumes were computed from these 3D surfaces. Numbers of pH3+ nuclei were quantified with the Spots detection function of Imaris (with background subtraction and mean spot diameters of 8 μm), and density of dividing cells were quantified as the number of pH3+ spots/total volume of structures. The measurement function in Imaris was used to annotate manually and quantify the major axis lengths (longest spine along anterior-posterior axis), minor axis lengths (diameter at the interface between anterior and posterior regions for elongated structures, based on anterior spherical shapes, and the longest diameter for non-elongated structures) and aspect ratios (defined as major axis lengths/minor axis lengths) per structure and condition. Data was exported from Imaris and analyzed in Python (version 3.9.18).

For spatial measurements along the A-P axis in fixed samples, and measurements of volumes, pH3 spots and pH3 densities along this axis, the same analysis steps were followed using python and napari (z-corrections, segmenting surfaces and detecting pH3+ spots). Images were rescaled to isotropic voxel sizes of 1 μm, corrected for the RI mismatch, and intensities were normalized based on the DAPI channel using a previously described pipeline^53^ . Gaussian blurring (kernel size 3,3,3) of normalized images and Otsu’s thresholding followed for surface segmentations, followed by binary dilation and closing operations. Then, the SpinePy framework^20^ was used to detect spines (defining an A-P axis, with A and P based on the (posterior) Bra domain) of structures, and to divide them in five equidistant segments along the A-P axis for 3D spatial measurements of pH3+ densities. For brightfield measurements of 2D shape changes during elongation, we applied a published pipeline^18^ to measure the areas, major axis and minor axis of elongating structures over time.

For measurements of cell division angles, stainings of pH3+ nuclei with clear chromosome arrangement in metaphase or in anaphases were used to define the mitotic plane, and to manually measure the angle of cell divisions to the local A-P axis (as defined by Bra staining), using the *Angle tool* in Fiji. Cell division angles to the A-P axis (*α*) were measured in posterior (Bra+), middle, and anterior regions, and were converted to the range of 0°-90°, by applying 180° - *α*, if *α* > 90°. Data was plotted combining angles in all regions for each time point.

For measurements of pH3 densities in fixed, stained and cleared mouse embryos at E8.5, a marching cubes measurement approach was done in Imaris. Cubes of 100×100×100 μm were used to measure the number of pH3+ spots in the cube volumes in the tailbud, PSM and somite tissues, with the same spot detection parameters as done for gastruloids.

### Analysis of surface actin intensity profiles

Quantifications of the posterior actin cap signal in fixed and stained gastruloids was performed based on Phalloidin (F-Actin) staining signals. Using Fiji, first a subset of 20-40 z-planes along the middle of structures was taken from each z-stack of volumetric imaging of actin. From this subset, a Sum projection of intensities was made, and a segmented line ROI with a 10 point of width was drawn along the gastruloid edge from the middle of the posterior end to the middle of the anterior end, in order to measure the intensity of surface actin along the A-P axis surface. Measurements were performed bilaterally, on each surface side from posterior to anterior, in each gastruloid. Raw intensities were exported from Fiji and analyzed in python. We first normalized the intensity of each individual curve by the average intensity of the first 50 μm from the posterior end. Then we obtained mean intensity curves and s.e.m. by performing ensemble averaging of normalized curves. The mean normalized intensity curves were fitted with the functional form 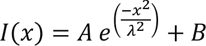, where A, B and *λ* are parameters. From the fitted gaussian exponentials, we obtained the lengthscale *λ* of the actin intensity decay from posterior to anterior. From the residuals of the fitted curves, a covariance matrix for the different fitted parameters (A, B, *λ*) and a 95% confidence interval (CI) were used to obtain the error in the actin lenghtscale. The same procedure was followed for the analysis of surface actin intensity decays in 72h human gastruloids. For mouse embryos fixed, stained and cleared at E8.5, the actin intensity at the tissue surface along the tailbud-PSM-somite regions was measured from reslices along the tailbud-somite curved axis of laterally or ventrally mounted embryos, in order to achieve an in-plane view of the actin signal across tissues even when embryos appeared curved. Measurements of mouse embryo surface actin intensity were performed from single resliced planes, and analyzed the same way as in mouse and human gastruloids.

### Droplet preparations, calibrations and injections in gastruloids and embryos

To measure mechanical stresses, fluorocarbon oil droplets were prepared similarly as previously described^23,24,26^ and adapted for measurements in gastruloids. HFE7700 fluorocarbon oil (3M Novec) was mixed with a final concentration of 4% (w/w) of fluorinated surfactant Krytox-PEG(600) (008-fluorosurfactant, RAN Biotechnologies) and 100 μM of a custom-made fluorinated Cy5 dye^54^ (FCy5). An additional 8% of a custom-synthesized fluorosurfactant^24^ (PEG_5_PFPE_1_) was added to lower the interfacial tension of the droplet to levels that enabled quantitative measurements of droplet deformations and stresses. Oil droplet mixtures were calibrated using a spinning drop tensiometer (KRÜSS), in order to determine their interfacial tension, as described before^24^. Calibrations were done in Ndiff medium and at 37°C, acquiring an interfacial tension *γ* (IFT) value of *γ* = 1.31 ± 0.02 mN/m for the oil droplet mixtures used for gastruloids. Injection of droplets into gastruloids and embryos was done with a pressure-controlled microinjector (Warner Instruments PLI-100A) set up in a laminar flow hood, adapting the injection time and pressure to produce droplets of 30-40 μm. Injections were performed in custom-made agarose pads with microwells to fit gastruloids and embryos. After injections, gastruloids and embryos were allowed to recover in an incubator at 37°C and 5% CO_2_ for 2-4h.

### Stress measurements with oil droplets

To measure stresses with oil microdroplets, we used similar methods as previously described^23^ , which rely on reconstructing 3D images of the droplets and obtaining their deformations. Imaging of oil droplets in mouse gastruloids, human gastruloids and mouse embryos was done with a Laser Scanning Confocal Microscope (Zeiss LSM 980) using a 40X\1,1 C-Apochromat water immersion objective (Zeiss), acquiring 1024×1024 pixels/frame with optimum z-step sizes (0.3-0.4 μm), in order to image droplets in maximum resolution z-stacks. Imaging was done with 2-3X digital zoom into droplets, with fastest dwelling time possible and using 2-line steps. 3-5 replicates the 3D scans of each droplet were done at 5 min intervals. Large view imaging of droplets for tissue context were also acquired with a 5X\0,25 objective (Zeiss) for each sample. 3D images of oil droplets were analyzed with an analysis software that we developed previously^25^ (Napari-STRESS v0.4.2). Droplet images were rescaled to obtain isotropic voxel sizes and blurred with sigma 1.0 before processing the Napari-STRESS software. To obtain the stresses from the surface deformations in napari-STRESS, we used the measured value of the interfacial tension. Samples with bad point cloud reconstructions (Gauss-Bonnett relative errors higher than 0.1) were excluded from analysis, since stresses cannot be properly computed in such cases. To display tissue anisotropic stresses *σ* on the ellipsoidal mode of the droplet (Fig. 3c), we used the expression: *σ* = 2*γ* (*H_e_* − *H_e,M_*), where *H_e_* and *H_e,M_* are the local mean curvature of the ellipsoid and the maximal mean curvature, respectively. To obtain the angles of droplets with respect to the anterior-posterior axis in mouse gastruloids, we measured the direction of the droplet’s main principal axis from the Napari-STRESS analysis and the local direction of the AP axis closest to the droplet from large view images for each sample. Angles between the main axis of droplets and of gastruloids were converted to the range of 0°-90° as explained before. For human gastruloids, droplet angles to the AP axis were measured manually in Fiji with the *Angle tool*.

### Laser ablation of mouse gastruloids

Mouse gastruloids generated from Lifeact-mScarlet mESC line for labelling of actin were used for two photon laser ablations of surface actin at anterior and posterior regions between 115-120h after aggregation. Samples were imaged using a Zeiss LSM980 confocal microscope with a 40X\1,1 C-Apochromat water immersion objective (Zeiss). For each gastruloid, first a z-stack of the whole structure or of anterior and posterior regions was acquired with a 0,6X digital zoom with z-steps of 3μm, in order to image the live actin structure before ablations. Then, a zoom-in was done into each region and ablations were performed with a femtosecond laser (InSight STDS-AX, Spectra-Physics) by applying a laser treatment on either circular ROIs of 30μm of diameter at the outer-most z-plane of structures or on rectangular ROIs of 50μm at the anterior and posterior edges of structures in equatorial z-planes. Ablation parameters used were: laser wavelength = 920 nm, laser power of 500-600 mW, and 3-6 repetitions depending on the region and depth in structures, imaging the ablation plane every 0.68 seconds before and after ablation for up to 80 frames. Analysis of surface rectangular ROI ablations was done in Fiji, by generating kymographs of tissue deformations with the *Reslice* plugin, and measuring tissue deformation over time after ablation at anterior and posterior regions based on the kymographs. Analysis of circular surface ablations was done in Fiji with the PIV Analyser plugin for analysis of 2D tissue movement after ablation at anterior and posterior regions, and measurements of the tissue recoil speed after ablation based on the velocities measured from PIV vectors.

### Perturbations with small drug inhibitors

Cell division perturbations in gastruloids and embryos were performed with Genistein (G2/M phase transition inhibitor, preventing cell entry into mitosis, Sigma G6649^21,55^), using a range of 1-100 μM for titration in mouse gastruloids, 10 μM for the wash-out experiment (keeping the drug for 6h from 96-102h, then washing two times with fresh pre-incubated NDiff medium, and allowing recovery until 120h) and 100 μM for treatment of human gastruloids and mouse embryos (since these structure sizes are bigger than mouse gastruloids). Additional cell division inhibitors used were Olomoucine II 7 μM (CDK inhibitor, Abcam ab141238^56^ ), Hydroxyurea 20 μM (S phase inhibition, Sigma H8627^57^) and Aphidicollin 2 mg/ml (S phase inhibition, Cell Signaling Technology 32774S^58^). For inhibitions of actin polymerization and activity, Cytochalasin D 5 μM was used (Sigma C8273^59^), and for inhibition of myosin activity Blebbistatin 10 μM (Cayman Chemical, Cay24171^60^). Samples were collected for fixation and staining after 24h of treatment for gastruloids (96-120h for mouse gastruloids; 48-72h for human gastruloids) and after 6h of treatment for mouse embryo tailbud explants. Samples with fresh pre-incubated Ndiff medium without inhibitors were used as controls.

### Statistics and reproducibility

Sample sizes are reported in figure legends. No statistical methods were used to predetermine sample sizes. Data visualization and analysis was performed in python using mean dot plots or line plots with s.e.m. error bars or shadows. For statistical testing between conditions, first Shapiro-Wilk tests were used to assess normality of distributions, and Levene’s test for variances. Paired data were tested with paired t-test if distributions were normal, or with Wilcoxon signed-rank tests otherwise. Independent two-group comparison were done with Student’s t-test when distributions followed normality and had equal variances, Welch’s t-test when normal but with unequal variances, or Mann-Whitney U test when non-normal. Experiments were not randomized and investigators were not blinded to allocation during experiments.

## Data Availability

Source data supporting these findings are available upon request.

## Code Availability

The SpinePY^20^ code used to analyze the gastruloid shapes in this article can be found at: https://gist.github.com/Cryaaa/010057456b3be0f868f2c077e1a44b04.

The napari-STRESS^25^ code to analyze the mechanical stresses with droplets can be found at: https://github.com/campaslab/napari-stress.

## Notes

### Competing Interest Statement

The authors have declared no competing interest.

